# *In Vitro/In Vivo* Assessment of Aripiprazole-Loaded Thiolated Arabinoxylan based Nanoparticles: An Innovative Approach for Targeted Schizophrenia Therapy

**DOI:** 10.1101/2024.02.13.580125

**Authors:** Mehwish Sikander, Ume Ruqia Tulain, Nadia Shamshad Malik, Arshad Mahmood, Alia Erum, Muhammad Tariq Khan, Asif Safdar

## Abstract

This study was conducted with the primary objective of improving the bioavailability of aripiprazole (APZ) through the development of nanoparticles using thiolated arabinoxylan (TAX) sourced from corn husk. TAX was synthesized via thiolation, employing thiourea as a thiol donor and hydrochloric acid as a catalyst. Characterization of TAX revealed a surface free thiol group content of 37.461 mmol/g, accompanied by an angle of repose measuring 0.393±0.035. Bulk density, tapped density, Hausner ratio, and Carr index fell within prescribed limits. Subsequently, APZ-loaded thiolated arabinoxylan based nanoparticles were fabricated using the ionotropic gelation method, with barium chloride serving as a cross-linker. Encapsulation efficiency was highest for formulation F4, at 97.1%±2.36. In vitro drug release demonstrated sustained release profiles at both pH 1.2 and pH 6.8, with F4 exhibiting the most favourable release kinetics. In vitro, characterization indicated that the optimized thiolated arabinoxylan based nanoparticle formulation had an average particle size of 211.1 nm with a Polydispersity Index (PDI) of 0.092 and a zeta potential of 0.621 mV. SEM imaging showed uniform, slightly spherical particles with minimal pores. DSC and TGA confirmed the conversion of APZ to amorphous states within the nanoparticles, enhancing solubility. Ex-vivo permeation studies exhibited favourable drug permeation. An In-vivo pharmacodynamics studies in a ketamine-induced schizophrenia rat model indicated the effectiveness of APZ loaded thiolated arabinoxylan based nanoparticles in behavioural tests, with no significant cataplectic effects observed. Acute oral toxicity assessments demonstrated the safety, with no mortality, no significant alterations in food and water consumption, or any histopathological abnormalities. In conclusion, these developed APZ-loaded thiolated arabinoxylan based nanoparticles hold promise for the effective treatment of schizophrenia without inducing toxic effects, showcasing their potential for clinical applications.

## Introduction

Pharmaceutical research strives to overcome a significant challenge i.e., enhancing the solubility and bioavailability of drugs, especially those plagued by low solubility and bioavailability. Nanotechnology has emerged as a transformative tool in this endeavour, presenting innovative solutions for drug delivery systems, notably in terms of pharmacokinetic optimization [1].

Nanocarriers have revolutionized drug solubility, offering a diverse range of options, including nanocrystals, nanosuspensions, liposomes, polymeric nanoparticles, and amorphous nanoparticles. These nanocarriers effectively augment the bioavailability of poorly soluble drugs. They hold the promise of resolving long-standing challenges, thereby improving drug efficacy, patient tolerance, specificity, and therapeutic ratios. Consequently, they have become the preferred choice over traditional medicines due to their ability to provide targeted drug delivery, controlled release, and enhanced therapeutic outcomes [2]. Central to the success of nanoparticle-based drug delivery systems is the judicious selection of suitable polymers. The characteristic particle size of these nanoparticles, typically ranging from 10 to 1000 nm, contributes to their remarkable performance. Natural polymers have gained recognition for their ability to enhance the therapeutic potential of both water-soluble and water-insoluble drugs, bolstering solubility, bioavailability, and retention within the body [3].

Arabinoxylan, sourced from materials such as corn husks, has emerged as a valuable polymer in these drug delivery systems, contributing to enhanced pharmaceutical results [4]. Recent research endeavours have concentrated on the development of thiolated arabinoxylans, also known as thiomers. These thiomers, created through the introduction of thiol groups into non-thiolated arabinoxylan, have proven instrumental in elevating solubility, drug permeability, and adhesion time to gastrointestinal mucosal membranes. Thiomers represent a significant breakthrough in controlled drug delivery systems, offering the potential for targeted administration routes and enhanced control over drug release profiles [5].

This manuscript focuses on a specific application of thiolated arabinoxylan-based nanoparticles that encapsulate APZ, a class IV BCS (Biopharmaceutics Classification System) drug characterized by poor solubility and permeability [6]. The overarching goal is to address the solubility and bioavailability challenges associated with APZ, ultimately improving therapeutic outcomes for patients [7].

Schizophrenia, a chronic and debilitating mental disorder that afflicts approximately 1% of the global population, necessitates continuous medication throughout an individual’s lifetime. Non-adherence to prescribed medications, particularly in the context of schizophrenia management, is a significant factor contributing to poor patient prognosis [8]. A complex interplay of factors, including genetic predisposition, environmental influences, substance use, brain chemistry, inflammation, and autoimmune diseases, has been postulated as risk factors in the development of schizophrenia. This debilitating disorder typically emerges at a young age, often before 18 years, and manifests a spectrum of symptoms, including positive, negative, and cognitive aspects. Managing schizophrenia remains a formidable challenge [9].

In the context of this study, we develop into the development of APZ-loaded TAX (Thiolated Arabinoxylan) based nanoparticles. These nanoparticles are synthesized using ionotropic gelation, facilitated by the cross-linking agent barium chloride. The primary objective of developing thiolated arabinoxylan-based nanoparticles is to enhance the solubility and bioavailability of APZ. This research encompasses a comprehensive analysis of the in vitro characteristics and in vivo behaviour of these nanoparticles, shedding light on their potential application as a drug delivery system for improved therapeutic interventions. These insights could contribute significantly to the management of schizophrenia and other challenging conditions.

The rationale for this study is deeply rooted in the need for innovative solutions to address the longstanding challenges associated with drug solubility and bioavailability. Poorly soluble drugs, like APZ, require sophisticated formulations and delivery systems to maximize their therapeutic efficacy. The quest for novel drug delivery technologies has led to the exploration of nanotechnology, a field that holds great promise in enhancing drug performance. Nanoparticles, particularly those based on thiolated arabinoxylan, have shown potential for improving drug solubility and bioavailability. This study recognizes the significance of developing nanoparticles to encapsulate APZ, a drug characterized by its low solubility and poor permeability. The choice of thiolated arabinoxylan as a polymer matrix for these nanoparticles is strategic, given its known benefits in drug delivery systems. Thiolation of arabinoxylan enhances the therapeutic potential of these nanoparticles [10]. Furthermore, the specific focus on APZ is driven by the pressing clinical need to improve the management of schizophrenia. Schizophrenia is a complex mental disorder that demands consistent and effective medication throughout a patient’s lifetime. Non-adherence to treatment regimens is a major concern in schizophrenia management and is associated with poor patient outcomes. Enhancing the solubility and bioavailability of APZ can contribute to more reliable and effective therapeutic interventions for individuals with schizophrenia, addressing a critical aspect of their care [11].

This study aims to bridge the gap between innovative nanotechnology and the practical needs of patients with schizophrenia. By developing and evaluating thiolated arabinoxylan-based nanoparticles loaded with APZ, we seek to offer a solution that could significantly impact the treatment of this challenging mental disorder. The potential advantages of enhanced solubility, bioavailability, and targeted drug delivery may hold the key to improved patient outcomes and adherence to treatment regimens, ultimately enhancing their quality of life and prognosis.

## Materials and methods

## Materials

The corn husks used in this study were procured from the local market in Chakwal. Various chemicals and reagents were employed, and their sources are as follows: Potassium dihydrogen phosphate and Disodium dihydrogen phosphate were obtained from Merck, Darmstadt, Germany, while glacial acetic acid was sourced from Merck, Germany. Hydrochloric acid (HCl) and sodium chloride (NaCl) were acquired from Supelco, USA. Methanol and ethanol were supplied by Supelco, USA, and acetonitrile was procured from Sigma-Aldrich, USA. Thiourea was obtained from Merck, Darmstadt, Germany. Aripiprazole (APZ) was generously provided as a gift by Genome Pharmaceuticals, Rawalpindi. Sodium hydroxide (NaOH) was sourced from Merck, Darmstadt, Germany. All chemicals and reagents used were of analytical grade.

## Methods

### First Method: Extraction of Arabinoxylan using autoclave

The process of extracting arabinoxylan from corn husk involved an alkaline extraction method, where the optimal concentration of sodium hydroxide (NaOH) was crucial for achieving a high yield. To do this, a 1% alkali solution was added to 10 grams of milled corn husk, maintaining a solid-to-liquid ratio of 1:10. The treated material was then exposed to steam in an autoclave at an elevated temperature of 121°C and 15 lbs pressure for 1h. The portion containing alkali-soluble arabinoxylan was separated by filtering it through a white plain muslin fabric. To adjust the pH of the arabinoxylan solution to 5.0, glacial acetic acid was added carefully. Corn husk arabinoxylan was obtained by mixing the arabinoxylan solution with ice-cold rectified alcohol in a 1:3 ratios. The resulting corn husk arabinoxylan was dried in an oven at a controlled temperature of 60±2°C. After drying, the arabinoxylan was weighed, ground using a grinder-mixer, and then stored at room temperature for future use [12]. Pictorial representation of stepwise arabinoxylan extraction is given in Figure 1.

**Figure 1.**
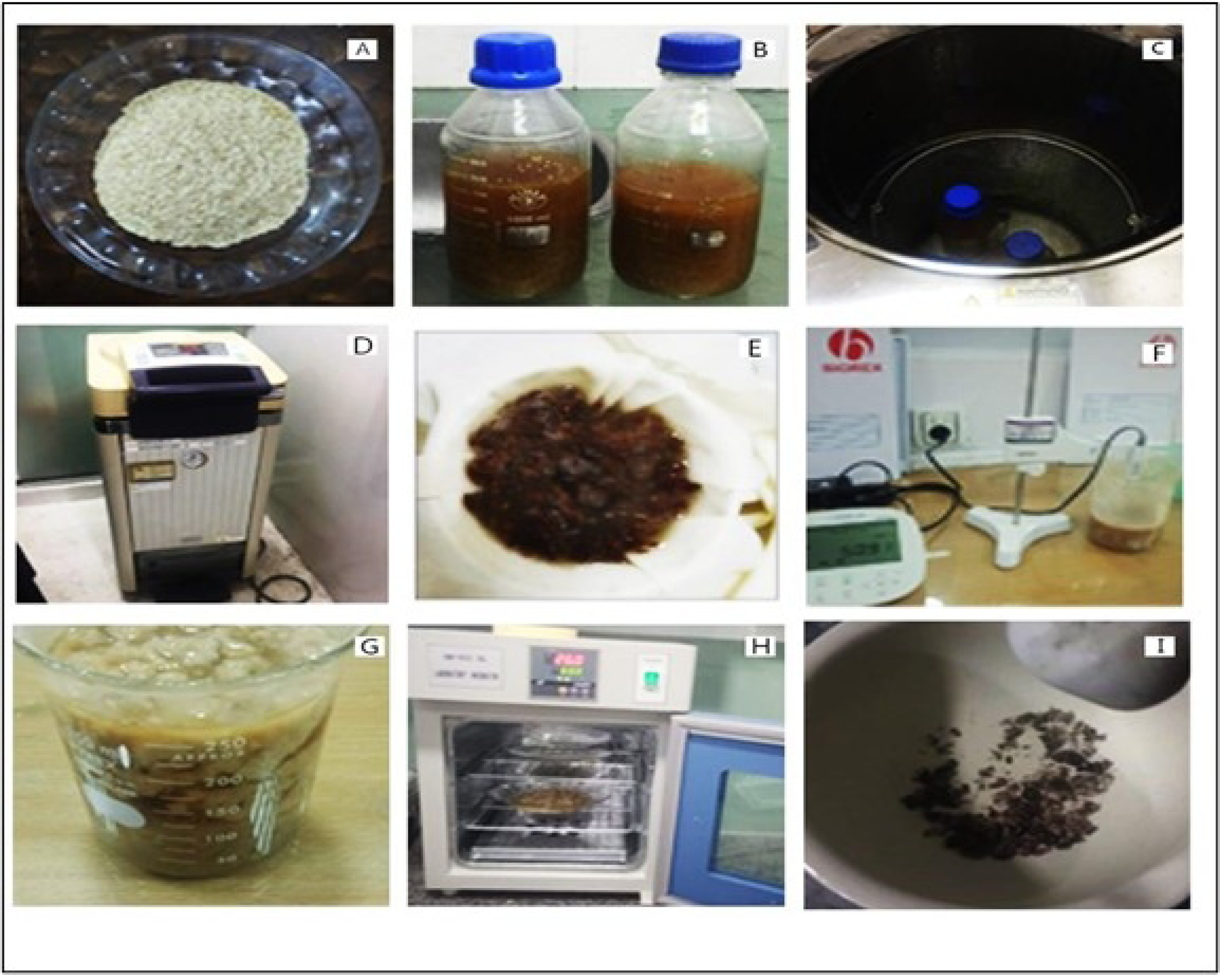
Pictorial representation of stepwise arabinoxylan extraction (A-I)

### Second Method: Alkaline Extraction of Arabinoxylan by centrifugation

To prepare the corn kernels, 800 g were combined with 2.4 L of water and brought to a boil at 100°C for one hour, with the addition of 8 g of slaked lime. The resulting cooking liquor, known as nejayote, was collected. The nixtamal produced was manually agitated with an additional 1000 mL of water, and this wash water was later added to the nejayote, adjusting the total volume to 2.8 L. Before extracting the nejayote, it was filtered under vacuum using plain white coarse unbleached cotton cloth with a pore size of 280 mm. This filtration step aimed to remove insoluble solid matter, leaving behind a fraction enriched with soluble solids. This fraction was then mixed with a 0.5 N solution of sodium hydroxide (NaOH) at a ratio of 3 g of solid to 45 mL of alkali solution. The mixture was kept at 25°C in darkness and placed on a rotatory shaker, agitating at a speed of 100 rpm for 8 h. Following agitation, the solution was centrifuged at 17,000 g at a temperature of 20°C for 15 minutes, causing particles to concentrate as pellets at the bottom of the centrifuge tubes. These centrifuged pellets were washed with water and subjected to another round of centrifugation (17,000 g at 20°C for 15 minutes). The supernatant and wash water from each centrifuged sample were combined, and 3N hydrochloric acid was added to adjust the pH to 4. These acidified supernatant samples underwent another round of centrifugation (17,000 g for 15 minutes at 20°C) and were subsequently filtered through Whatman filter paper with a pore size of 2.7 mm. To the obtained filtrate, 65% v/v ethanol was added, and it was left overnight at 4°C to induce precipitation. The precipitate was recovered through centrifugation (16,000 g at 4°C for 15 minutes) and subjected to air-drying to remove ethanol. The dried arabinoxylan was weighed and then ground using a grinder-mixer. Finally, the ground arabinoxylan was stored at room temperature for future use [13].

### Percentage yield

The calculation of the percent yield of obtained arabinoxylan can be expressed using Equation 1:

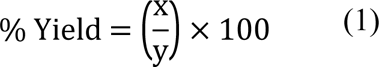

Where:

% Yield represents the percentage yield of arabinoxylan.

x represents the weight of arabinoxylan.

y represents the weight of corn husk [14].

### Thiolation of Polymer

The thiolation of arabinoxylan was performed using a well-established method, with hydrochloric acid serving as the catalyst for the reaction. Initially, a 1% aqueous solution of TAX was prepared by dissolving 1 g of the polymer in 100 mL of distilled water. This solution was continuously stirred using a magnetic hot plate stirrer at a speed of 500 rpm. Next, 2 g of thiourea were added and stirred for ten minutes. To initiate the reaction, a catalytic amount (4-5 drops) of HCl was introduced. The polymer solution was then placed in a water bath at 70°C and allowed to react for 90 minutes. After the allotted time, the polymer solution was removed from the water bath, and methanol was added to cool down the solution, resulting in the formation of precipitates. The resulting precipitate was carefully filtered and subjected to multiple washes with hydrochloric acid and distilled water to remove any unreacted thiourea. Following the washing steps, the final material was cooled to -80°C and subsequently lyophilized at -47°C and 0.013 mbar pressure to obtain a dried polymer. The freeze-dried polymer was stored in a sealed glass vial for future use [15]. Schematic representation of stepwise thiolation of arabinoxylan is shown in Figure 2.

**Figure 2.**
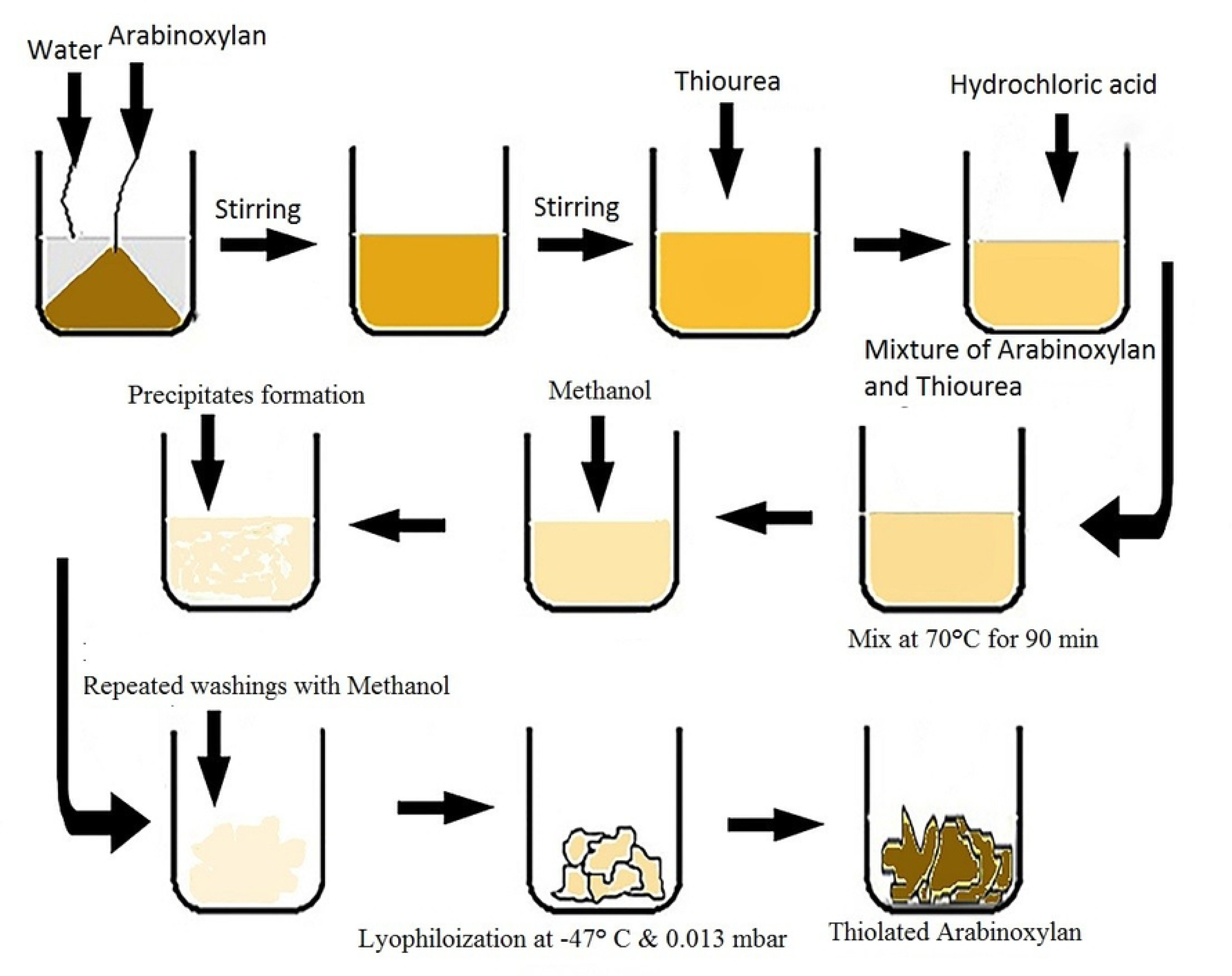
Schematic representation of stepwise thiolation of arabinoxylan

## Characterization of Thiolated arabinoxylan

### Quantification of Thiol substitution by Ellman’s method

The thiol content attached to the AX backbone was quantified using the Ellman’s reagent method. To establish a standard curve, a 0.2% w/v stock solution of thiourea (1 mL) was initially prepared by dissolving thiourea in 0.5 M phosphate buffer saline at pH 8. Serial dilutions were then created within the concentration range of 10 to 100 μg/mL. These dilutions were analysed spectrophotometrically at a wavelength of 280 nm. For the assessment of thiol content in AX, a 2% w/v solution of thiolated arabinoxylan was prepared by dissolving it in distilled water. Simultaneously, 150 µl of Ellman’s reagent was prepared by dissolving 3.96 mg of the reagent in 10 mL of 0.5 M phosphate buffer saline (pH 8). In a beaker, 150 µl of Ellman’s reagent, 1350 µl of PBS, and 150 µl of TAX were mixed in 1350 µl of deionized water. This mixture was incubated at room temperature for 10 minutes. Subsequently, the sample was analysed using a spectrophotometer to determine the absorbance at 412 nm. The calibration curve of thiourea was employed to calculate the free thiol groups present in arabinoxylan [16].

## Micromeritic Studies

### Bulk Density, Tapped Density, Carr’s Index, Hausner’s Ratio and Angle of Repose

Micromeritic studies were systematically undertaken to evaluate the rheological characteristics of the powdered samples. These assessments covered a spectrum of parameters, including bulk density, tapped density, Carr’s index, Hausner’s ratio, and the angle of repose, in accordance with established protocols [17].

Angle of repose quantifies the maximum angle attainable between the freestanding surface of a powder pile and the horizontal plane. It serves as a simple yet informative measure of particle resistance to movement, reflecting both internal cohesion and frictional effects within the powder. An angle of repose up to 40° indicates favorable flow, while angles exceeding 50° suggest poor or absent flow [18]. The angle of repose was determined by freely pouring arabinoxylan through an 8 mm diameter glass funnel onto a flat surface, forming a cone-shaped pile. Measurements of the pile’s height and diameter allowed the calculation of the angle of repose using the tangent formula [19]. The angle of repose (h) was determined through the utilization of Equation 2, and this experimental procedure was conducted in triplicate:

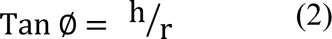

Where h is height and r is radius of powdered heap [14]

In adherence to USP Guidelines, bulk density was assessed by adding a weighed quantity of powder (passed through a 1 mm mesh sieve screen) into a 100 mL graduated cylinder with a 100 cm3 capacity. The powder was gently flattened without compression, and the apparent volume (Vb) was recorded [26]. Equation 3 was used to calculate the bulk density (ρb) and was measured in cm^3^:

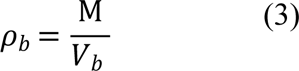

where M is mass of the polymer and *V*_*b*_ is its apparent volume.

Tapped density was determined according to USP Guidelines, with the powdered sample placed in a graduated cylinder. Mechanical tapping was applied initially for 500 times, followed by recording the tapped volume to the nearest graduated unit. An additional 750 taps were performed, and the difference between the two tapped volumes was assessed. If the difference remained below 2% for all powders, the second tapped volume (Vf) was considered the final tapped volume for tapped density calculations [27]. The formula used to calculate the tapped density (*ρ*_*t*_) in cm3 is mentioned in equation 4:

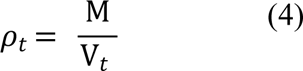

where M is mass of the polymer and *V*_*t*_ is tapped volume of powder [15].

The Carr’s Compressibility index, a measure of powder compressibility, was employed to assess the flow properties of the samples. A higher Compressibility index (C.I %) indicates greater powder compressibility. The formula used to calculate (C.I %) is mentioned in equation 5.

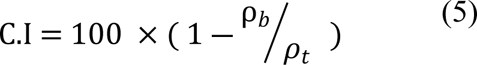

Where ρ_*b*_ is bulk density and *ρ*_*t*_ is tapped density of powder [16]

Hausner’s Ratio is a numerical value which is correlated to the flowability of powder [17]. The Hausner ratio is defined as the ratio between the tapped and bulk density of powders [18]. It is calculated by the formula:

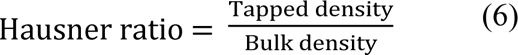

### Preparation of Thiolated Arabinoxylan based Nanoparticles

The ionotropic gelation method was used for the preparation of thiolated arabinoxylan based nanoparticles with slight modifications using Barium chloride as cross-linker. Solution A was prepared by dissolving the required amount of TAX as mentioned in Table 1 into the deionized water at a concentration of 5 mg / mL using gentle heat and continuous stirring at hot plate magnetic stirrer at stirring rate of 500 rpm. Similarly solution B was prepared by dissolving accurately weighed APZ in 2% glacial acetic acid at a concentration of 1mg / 1mL at room temperature through continuous stirring to form a clear solution. Both solutions were mixed with each other followed by continuous stirring to get a homogeneous solution. Solution C was prepared by dissolving barium chloride concentration (as indicated in table) into the distilled water with continuous stirring at room temperature. The resultant solution was filled into the syringe and was added dropwise via 20 Gauge needle syringe into Barium chloride solution with constant stirring at room temperature. An opalescent suspension was obtained which was subjected to ultracentrifugation at 12,000 rpm for 30 minutes. Nanoparticles were formed and settled at bottom which were subjected to lyophilization at -50°C and were stored at 4°C for further analysis [19].

**Table 1.** Preparation of Thiolated arabinoxylan based nanoparticles by Ionotropic Gelation Method.

| TAX based nanoparticles code | Drug (APZ) | Polymer (TAX) | Cross-linker (BaCl<sub>2</sub>) |
| --- | --- | --- | --- |
| F1 | 10 mg | 20 mg | 5 % |
| F2 | 10 mg | 30 mg | 5 % |
| F3 | 10 mg | 40 mg | 5 % |
| F4 | 10 mg | 50 mg | 5 % |
| F5 | 10 mg | 20 mg | 2 % |
| F6 | 10 mg | 20 mg | 3 % |
| F7 | 10 mg | 20 mg | 4 % |

## Characterization of Aripiprazole TAX based nanoparticles

### Encapsulation Efficiency

The encapsulation efficiency of each APZ-loaded TAX-based nanoparticle was evaluated by carefully removing unentrapped drug in the supernatant solution. This was achieved through centrifugation of the nanoparticle suspension at 15,000 rpm for 30 minutes, resulting in the settling of the water-insoluble unentrapped drug as a pellet at the bottom of the centrifuge tube. The clear supernatant was then collected and filtered for clarity. To quantify the concentration of the unentrapped drug in the filtered supernatant, its absorbance at 259 nm was measured using a UV-Visible Spectrophotometer (UV 1700, Shimadzu, Japan) [19]. The encapsulation efficiency (% EE) was calculated using the formula as mentioned in equation 7:

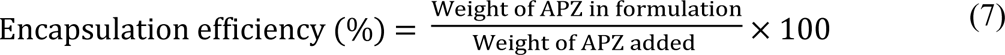

### In-Vitro Dissolution and Drug Release Kinetics

Aripiprazole release from thiolated arabinoxylan-based nanoparticles was assessed using the dialysis method, also referred to as the open-ended tube method. In this study, each thiolated nanoparticle, loaded with 10 mg of Aripiprazole, was suspended in 2 mL of two distinct dissolution media. These suspensions were enclosed within dialysis cellulose membrane bags with a molecular weight cutoff of 12,000 (D9402-100FT; Sigma-Aldrich, Steinheim, Germany), which were then sealed and placed into a dissolution basket. The in-vitro dissolution study was conducted using dissolution media with pH values of 1.2 (simulated gastric fluid) and 6.8 (simulated intestinal fluid), simulating physiological conditions, as part of the investigation involving TAX-based nanoparticles [21].

Release studies were conducted in a buffer containing 0.5% Sodium Lauryl Sulfate (SLS) to establish ideal sink conditions, as Aripiprazole exhibits limited solubility in standard buffer solutions [22]. The release assessments were performed under these sink conditions with a rotation rate of 75 rpm at a controlled temperature of 37 ± 0.5°C. The study involved two-time frames: a 2 h examination in Simulated Gastric Fluid (SGF) and a 48-h investigation in Simulated Intestinal Fluid (SIF). At defined intervals of 1, 2, 3, 4, 5, 6, 8, 12, 24, and 48 h, 2 mL aliquots were withdrawn from the samples, and the same volume of fresh dissolution medium was replenished accordingly. The collected sample aliquots were subsequently filtered and subjected to analysis using a UV-Visible spectrophotometer, specifically at 259 nm, to determine the Aripiprazole content [23].

Aripiprazole (APZ) loaded TAX-based nanoparticles were subjected to in-vitro drug release analysis using various kinetic models, facilitated by the DD Solver, a Microsoft add-in tool. Specifically, the release profiles were evaluated using Zero order, First order, Higuchi, Korsmeyer-Peppas, and Hixson-Crowell models [24-29].

## Characterization of Optimized TAX based nanoparticles

### Particle Size, Zeta Potential and Polydispersity Index

The mean particle size, polydispersity, and zeta potential of the TAX-based nanoparticles were determined using a Zetasizer Nano ZS instrument (Malvern Instruments Ltd., Worcestershire, UK). Photon correlation spectroscopy was employed to measure the mean size and polydispersity. The polydispersity index (PDI) values, ranging from 0 to 1, indicated the degree of size distribution uniformity in the TAX-based nanoparticles, with higher values signifying a less homogeneous distribution [30]. To analyse these parameters, the developed TAX-based nanoparticles were diluted fivefold using deionized water previously filtered through a 0.45 μm membrane filter. This analysis was conducted in triplicate to ensure consistency and reliability [31].

### Fourier Transform Infrared Spectroscopy (FTIR)

The FTIR spectrum reveals absorption peaks corresponding to the vibrational frequencies of atomic bonds in the polymer, making it a valuable tool for qualitative analysis. The intensity of these peaks offers information about the nature of the materials present [32]. In this study, FTIR was used to analyse arabinoxylan and TAX employing an FTIR spectrophotometer from Bruker (Germany) through the KBr pellet method. This method generates spectra by assessing electrostatic interactions among the particles [33]. To prepare the samples, the polymer was ground into a powdered form and mixed with potassium bromide (KBr) at a 1:100 ratio (sample to KBr). This mixture was then subjected to a hydraulic press with a pressure of 5 tons for 5 minutes to form pellets. Thirty scans of these pellets were collected at a resolution of 2 cm^-1^, spanning the range from 4000 to 400 cm^-1^ [34]. By examining characteristic and identifiable peaks in the sample, any unusual peaks observed in physical mixtures were also evaluated [35].

### Scanning Electron Microscopy (SEM)

The structural morphology of TAX-based nanoparticles was investigated using a scanning electron microscope (SEM) - JSM-7610 F from JEOL - operating at 25 keV. To conduct this analysis, 1 mg of the nanoparticles was placed onto a double-sided carbon adhesive disk with a diameter of 12 mm. The disk was then secured for microscopic examination and inserted into the sample compartment of the SEM.

The imaging process took place in a low-vacuum SEM environment with a pressure of 10 Pascal in the sample compartment. The SEM instrument was equipped with an electron gun with an accelerating voltage of 25 kV and a filament current of 60 µA, allowing for high-resolution imaging and detailed analysis of the nanoparticles’ structural features [36].

### Differential Scanning Calorimeter (DSC) and Thermogravimetric Analysis (TGA)

Differential Scanning Calorimetry (DSC) is employed to investigate the thermal properties of various substances, including Aripiprazole (APZ), thiolated arabinoxylan, a physical mixture of APZ and thiolated arabinoxylan, and Thiolated Arabinoxylan-based nanoparticles. The measurements were conducted using a Thermal analyser (DSC Q2000 V24, Seoul, Korea) that was calibrated using highly pure Indium and operated under a constant nitrogen flow with a flow rate of 50 mL/min. During the analysis, heat was supplied to the samples at a rate of 20°C/min. In the first run, the samples were heated to a temperature of 200°C, and subsequently, the melted samples were rapidly cooled to -20°C. In the second run, the heat supplied to the samples ranged from -20 to 200°C, and changes in heat capacity in the amorphous sample from the second heating run were determined [37]. This DSC test aimed to provide insights into the thermal behaviour of APZ, thiolated arabinoxylan, their physical mixture, and the TAX-based nanoparticles.

Furthermore, the study also involved Thermal Gravimetric Analysis (TGA) and was conducted using a TGA/DSC-1 instrument from Mettler Toledo, USA. The heating rate was set at 10°C/minute, and the analysis was carried out under a nitrogen atmosphere with a flow rate of 40 mL/min [38].

### Powder X-ray diffraction (PXRD)

X-ray Diffraction (XRD) is an analytical technique employed to study the characteristics and nature of the prepared nanoparticles [39]. The X-ray diffraction patterns of the arabinoxylan samples under study were acquired using an X-ray diffractometer from Philips X-ray Analytical, based in Amsterdam, The Netherlands. For the analysis, a precisely measured quantity of the sample was loaded into the sample holder’s cavity and then compressed. Subsequently, a glass slide was used to ensure the sample’s smoothness. The samples were scanned within a diffraction angle (2*θ*) range from 0° to 80° using a nickel-filtered Cu-K*α* radiation source. The XRD instrument operated at a voltage of 35 kV and a current of 25 mA. It featured a scanning speed of 0.05 min⁻¹ and had a division slit of 1.25° and a receiving slit of 0.3 mm [40].

### Ex-vivo intestinal Permeation Study

The permeation assessment of Aripiprazole (APZ) from thiolated nanoparticles was conducted employing the everted intestinal sac method with slight modifications. To evaluate drug permeation, the study was carried out in simulated gastric fluid (SGF) at pH 1.2 and simulated intestinal fluid (SIF) at pH 6.8 for a duration of 24 h. Ethical approval for all experiments involving experimental rats was obtained from the Institutional Animal Ethics Committee, specifically the Pharmacy Research Ethics Committee (PREC) of the Capital University of Science and Technology (CUST) in Islamabad, Pakistan.

Male Wistar rats weighing between 200-250 g were humanely euthanized through cervical dislocation using excess ether as an aesthetic. The small intestine was swiftly isolated, and the colon segment of the small intestine was carefully collected and cleansed of blood and debris using a saline solution [41]. The isolated colon segment was then gently inverted by passing a glass rod through it. Following this, a long segment (4 cm) was securely knotted with a thread and a sac was cut open from the main colon length. A second ligature was loosely fastened around the open end of the sac, and a needle connected to a syringe was inserted. The loose ligature was tied over the needle, and a suspension of thiolated nanoparticles equivalent to 10 mg of APZ was introduced into the sac [42].

The sacs were placed in separate conical flasks, each containing different media: 200 mL of SGF and 100 mL of SIF. The media were continuously aerated with a mixture of 95% O2 and 5% CO2. The conical flasks were maintained in a thermostatic water bath at 37 ± 0.5 °C. At regular intervals, 2 mL samples were withdrawn and replaced with an equal volume of the same medium. These samples were subsequently analysed for permeated drug content at 259 nm using a UV spectrophotometer [43].

### *In-Vivo* Pharmacodynamic studies

Pharmacodynamic studies of the optimized TAX-based nanoparticle formulation were carried out using Ketamine to induce schizophrenia in rats. Repeated dosing of Ketamine replicated positive, negative, and cognitive symptoms. Following acute Ketamine administration, abnormal expression of mTOR was observed in brain tissues, suggesting potential association with neurological disorders, including schizophrenia [44].

### Animals

For this study, male Wistar rats were acquired from Capital University of Science and Technology, Islamabad, with an initial weight ranging from 250 to 260 grams. These rats were group-housed, with four rats in each cage, within a carefully controlled laboratory setting. The laboratory maintained a consistent 12:12 light/dark cycle, with lights on at 06:00 and off at 19:00. The ambient temperature was consistently maintained at 21 ± 1°C, and the humidity level was kept at 55±5%.

The rats had unrestricted access to both food and tap water, allowing them to feed and drink as desired. Experimental procedures were conducted during the light phase, specifically between 07:00 and 17:00. To ensure the rats were acclimated to the laboratory conditions, a five-day adaptation period was implemented before the initiation of the experiments.

### Experimental Design

The rats were stratified into four groups, each consisting of four individuals:

Group I: Rats were administered the vehicle exclusively, consisting of normal saline delivered orally in a manner that paralleled the regimen.

Group II: Rats were subjected solely to Ketamine at a dose of 50 mg/kg, administered intraperitoneally, for a continuous period of 14 days. The 50 mg/kg Ketamine dose was prepared by withdrawing 0.25 mL of a 50 mg/mL Ketamine solution and administering this 0.25 mL dose to each rat once daily.

Group III: Rats were administered an APZ suspension, containing Aripiprazole at a concentration of 3 mg/kg, once daily, with a 30-minute interval preceding the administration of Ketamine at a dose of 50 mg/kg, intraperitoneally. APZ, at a dose of 1 mg/kg, was dissolved in 1 mL of normal saline (0.9% w/v) and administered orally via gastric gavage. Group IV: Rats received APZ nanoparticles at a dose of 3 mg/kg, administered 30 minutes before the administration of Ketamine at a dose of 50 mg/kg, intraperitoneally.

All drug administrations were conducted daily for a duration of 14 days, and it is noteworthy that all drug solutions provided to the rats were freshly prepared [45-49].

### Doses Selection

Rats were administered a daily dose of APZ at 3 mg/kg via oral route [50-51], in addition to ketamine at 50 mg/kg via intraperitoneal injection per day. Given the relatively shorter half-life of APZ, it was administered three times a day, with 8 h intervals between each administration. This dosing regimen aimed to achieve a concentration like that observed in humans when APZ is administered orally once per day [52].

### Behavioural Tests

The primary symptoms of psychosis in this study encompassed behaviour patterns, which were characterized by tests such as catelepsy, passive avoidance test, forced swim test, and maze test. These behaviours encompassed a range of symptoms, including positive or bizarre symptoms, negative or depressive symptoms, and cognitive symptoms. The behavioural parameters discussed below were closely associated with the signs and symptoms typically observed in schizophrenia [53].

### Catalepsy Study

This test was conducted to evaluate the impact of the developed APZ-TAX-based nanoparticles on extrapyramidal side effects associated with APZ. Animals were subjected to the catatonia test [54]. In this experiment, rats were gently held by their tails, and their forepaws were placed on a horizontal bar until the rats could grasp it. The bar was positioned at a height of 9 cm, while the hind paws, along with their tails, remained relaxed on the floor. Readings were taken for each animal at 1, 4, 8, and 24 hours after the administration of the ARP-TAX-based nanoparticles. The duration during which the rats maintained their forelimbs on the bar was recorded, with a maximum time limit of 30 seconds. Rats that held onto the bar for 30 seconds or more were considered to exhibit catalepsy [55].

A maximum of three attempts were made for each rat to remain on the rod for 30 seconds, and testing was concluded once the rats achieved the predetermined criterion time. The mean duration from these trials for each rat was subsequently calculated [56,57].

### Forced Swimming Test

This test serves as a highly reliable method for rapidly screening schizophrenics drugs to assess their efficacy in reducing immobility time in rats, indicative of their antischizophrenic effects. The initial 15 minutes served as a habituation period and were excluded from the analysis, while a 5-minute test period was conducted to minimize the impact of any drug-induced effects on memory. During the test, rats were placed in a transparent plastic container (measuring 47 cm in height and 38 cm in diameter) filled with 38 cm of water at a temperature of 25°C. The rats were compelled to swim in this inescapable environment. Initially, the animals made attempts to escape, but eventually, they displayed immobility. The entire duration of immobility, including passive swimming, was recorded. Passive swimming involved floating vertically in the water and making only the necessary movements to keep their heads above the water. After the test, the rats were removed from the water, dried with a towel, and returned to their cages. To maintain consistency, each experimental group of rats was assessed between 9 am and 11 am to minimize the potential influence of circadian variations on the test results [58].

### Tail Suspension Test

In the tail suspension test, each rat was suspended by its tail at a height of 50 cm using adhesive tape, and climb stoppers were employed to prevent the rat from grabbing onto its tail. The test was conducted for a duration of 5 minutes, during which the time spent struggling and the time spent immobile were recorded for each rat. Immobile behaviour was characterized by passive hanging and the absence of movement, except for whisker mobility and respiration. The period during which all movements, except for whisker mobility and respiration, were absent was defined as the immobility period [59].

### Passive-Avoidance Test

The impact of APZ on learning behavior and memory was evaluated through the implementation of the passive avoidance test. The test apparatus consisted of two compartments: a light compartment (measuring 200 × 250 × 200 mm) and a dark compartment (measuring 200 × 150 × 200 mm), both equipped with a grid floor system. These two compartments were separated by a sliding door. During the training phase, the rats were initially placed in the light compartment. After a 10-second interval, the door was opened, allowing the rats to enter the dark compartment. The time taken by each rat to enter the dark compartment was carefully recorded. Rats that took more than 100 seconds to cross into the dark compartment were excluded from the study. Once a rat entered the dark compartment, the door was promptly closed, and the rat was returned to its cage.

The acquisition trial was conducted 30 minutes following the habituation trial. Rats were gently placed in the light compartment, and after a brief 5-second period, the sliding door was opened. As the rats crossed into the dark compartment, the door was closed, and they received a mild foot-shock characterized by 50 Hz for 5 seconds at an intensity of 0.2 mA via the grid floor of the dark room, administered by a current shock generator. After a 20-second interval, the rats were removed from the dark compartment and temporarily placed in their home cage. After a 2-minute break, the test was repeated. The rats received a foot shock each time they entered the dark compartment with all four paws. If any rat failed to cross into the dark compartment and remained in the light compartment for 2 minutes, the training session was terminated. The evaluation included an assessment of the frequency of trials, specifically the number of entries into the dark compartment, with a maximum of three training sessions conducted for each rat [60].

### Elevated Plus-Maze Model

The Elevated Plus-Maze test is employed to assess fear and anxiety-related behaviour in rats, particularly in the context of schizophrenia research. This apparatus consists of a cross-shaped maze constructed from black plexiglass, comprising two open arms positioned opposite each other, each measuring 30.5 cm, and two closed arms similarly arranged, each measuring 30 x 20 cm. The maze includes a central area measuring 5.5 cm in width and is elevated to a height of 40 cm above the floor. The evaluation of anxiolytic effects involves monitoring the reduction of anxiety and an increase in entries into the open arms and exploration time. Parameters such as the frequency of entries into the arms and the time spent in both the closed and open arms are precisely measured. After each trial, the apparatus is thoroughly disinfected using a 70% alcohol solution to maintain cleanliness and hygiene during the experimental procedures [61].

### Acute Oral Toxicity Study

*In-vivo* acute oral toxicity studies were conducted using male Wistar rats weighing between 250-260 g, following the established protocols. The rats were subjected to daily observations over a 14-day period, with a focus on detecting potential signs such as signs of illness, treatment reactions, changes in mucous membranes, alterations in eye and skin color, fur conditions, behavioral changes, diarrhea, salivation, tremors, coma, or sleep. Furthermore, the body weights of the rats were recorded on Day-1 before the commencement of the experiment, as well as on Day-3, Day-7, and Day-14. Food and water consumption were also meticulously documented and compared with control data.

Upon the conclusion of the 14-day experimental period, blood samples were collected from the posterior vena cava of the heart through cardiac puncture while the rats were under chloroform anesthesia. These blood samples were divided into various vials for subsequent hematological and biochemical analyses, which encompassed assessments of liver profiles, renal profiles, and lipid profiles. The resulting data were subjected to statistical analysis using one-way ANOVA, with statistical significance indicated by a *p-value* of less than 0.05 [62].

## Results and Discussion

### Percentage Yield

The percentage yield of extracted arabinoxylan with reference to the total mass of the husk was calculated. The second extraction method yielded a higher percentage (42%± 0.65) of arabinoxylan compared to the first method (19% ± 0.87). The extracted arabinoxylan exhibited solubility in DMSO at a concentration of 1mg/10 mL at 40°C. However, it remained insoluble in various organic solvents, including ethyl alcohol, methyl alcohol, chloroform, acetonitrile, acetone, diethyl ether, hexane, formaldehyde, and glacial acetic acid.

## Characterization of Thiolated arabinoxylan

### Estimation of Thiol contents using Ellman’s Reagent and calibration curve

The thiolation process of arabinoxylan involved the reaction of arabinoxylan with thiourea. The quantification of thiol groups attached to the arabinoxylan backbone was determined using Ellman’s reagent method and analysed via a spectrophotometer. Ellman’s reagent, characterized by its hydrophilic nature, readily reacts with free thiol groups, resulting in the formation of a yellow-coloured product. A calibration curve of thiourea was established, as depicted in Figure 3. The thiol content of TAX was measured at 37.461 mmol/g of TAX. The detection of thiol groups provides compelling evidence for the successful thiolation process [20].

**Figure 3.**
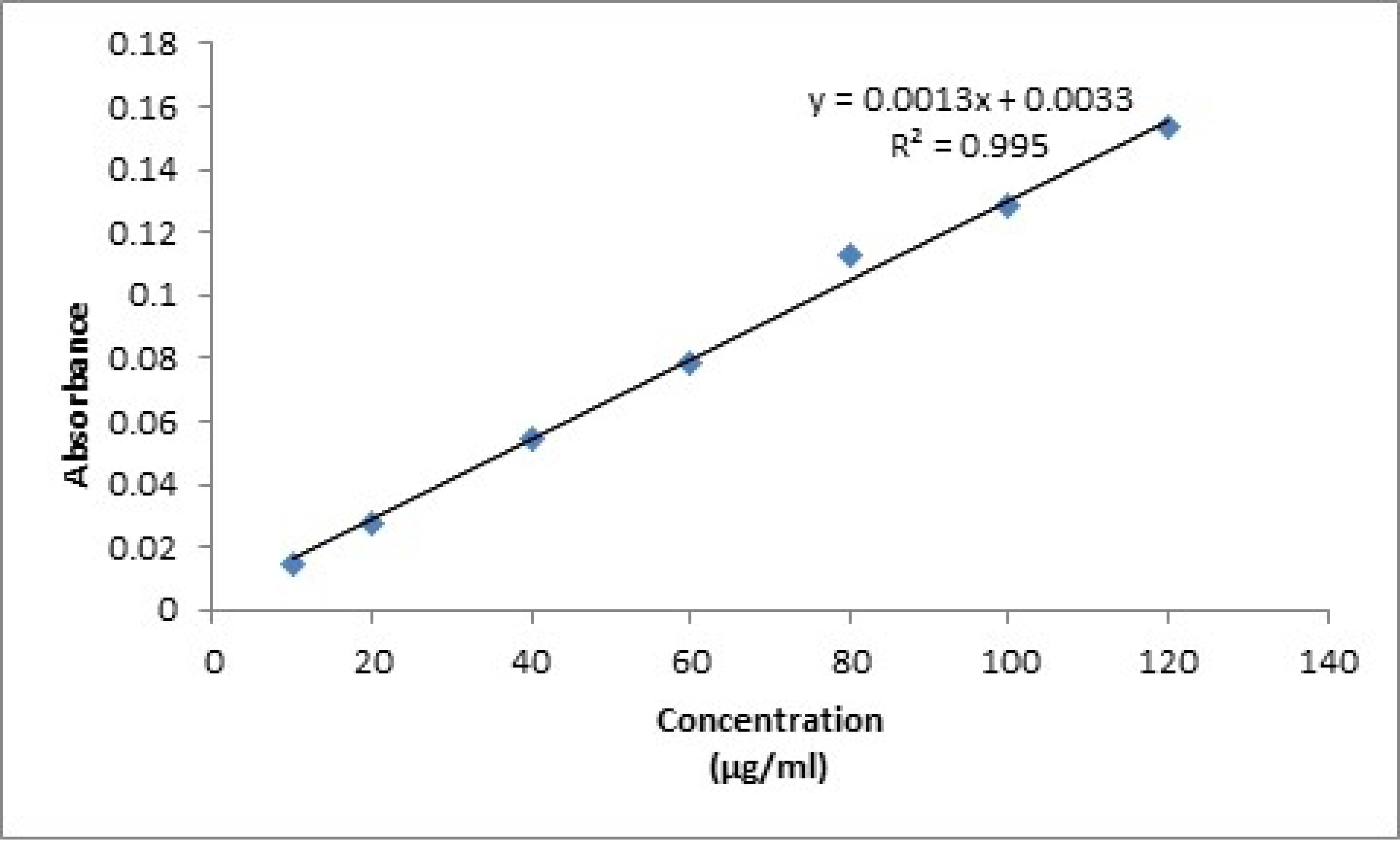
Quantification of thiol contents

### Micromeritics

The micromeritic characteristics of AX and TAX were examined to assess the flow properties of the polymers. Compared to AX, TAX exhibited favourable flow properties, Notably, TAX demonstrated an angle of repose of 0.393±0.035, well below the standard limit of 0.5, indicating excellent flow. The recorded bulk density and tapped density for TAX were 0.597±0.063 cm3/mL and 1.015±0.008 cm3/mL, respectively, falling within the accepted range. The Hausner ratio, at 1.441±0.017, and the Carr index, at 29.667±0.700, also met the standard limits, further emphasizing the excellent flow and good compressibility of the developed nanoparticles. These results align with the prescribed values, confirming the enhanced flow properties and compressibility of TAX in comparison to AX.

## Characterization of Aripiprazole TAX based nanoparticles

### Encapsulation Efficiency (EE)

The APZ-TAX NPs were synthesized using the ionic gelation method, employing varying quantities of polymer and cross-linkers to optimize the nanoformulation. Seven formulations (F1 to F7) were selected based on their encapsulation efficiency, with diverse formulation parameters explored. Table 2 outlines the composition and Encapsulation Efficiency (EE) of thiolated arabinoxylan-based nanoparticles (F1, F2, F3, F4, F5, F6, and F7). An effective drug delivery system should ideally exhibit a robust association with drugs. The EE for all TAX-based nanoparticles demonstrated a satisfactory level, indicating the presence of sufficient ionic interactions between aripiprazole and thiolated arabinoxylan. This interaction facilitated the efficient entrapment of the drug within the formed TAX-based nanoparticles [21]

**Table 2.**
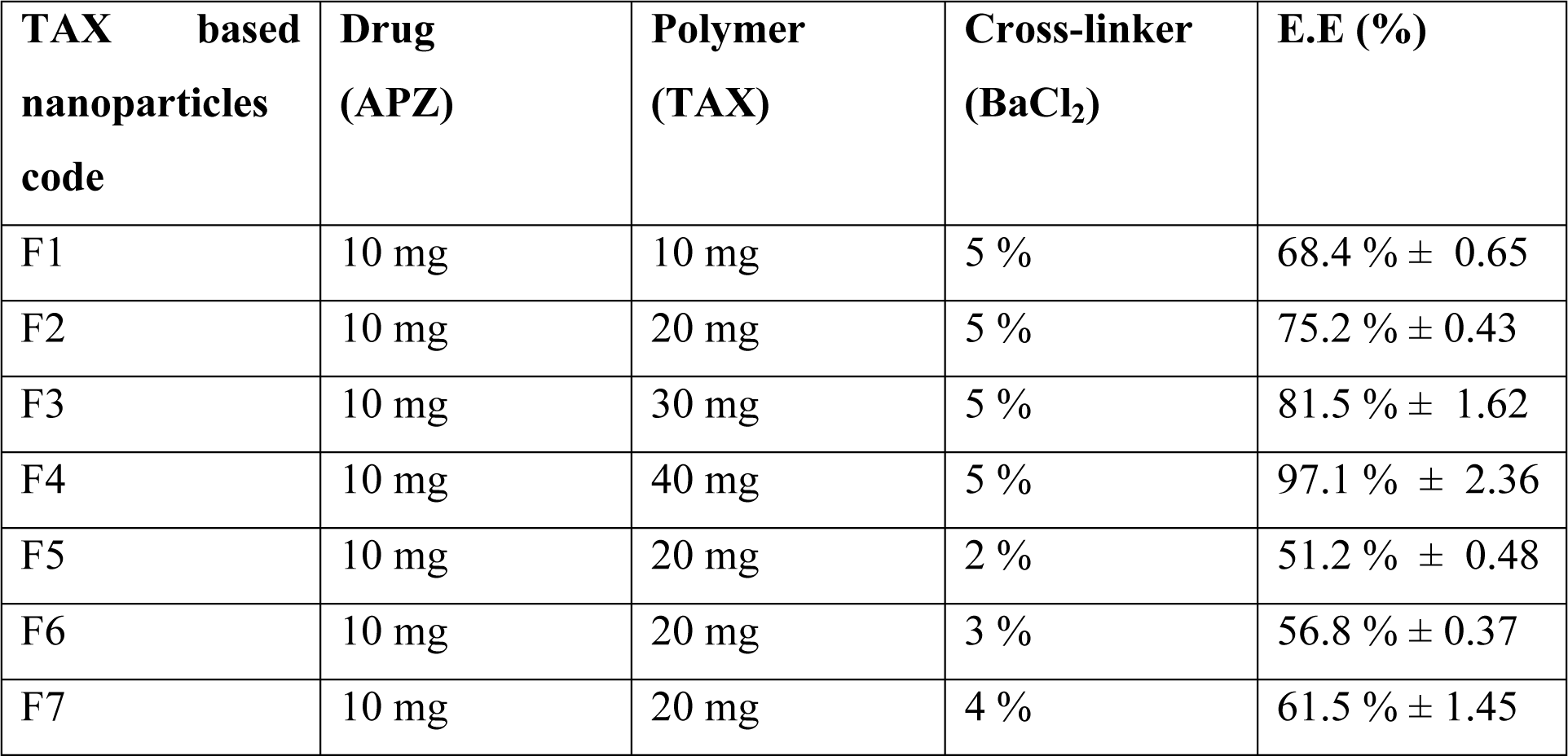
Encapsulation Efficiency of Thiolated arabinoxylan based nanoparticles (Mean ± SD, n=3)

| TAX based nanoparticles code | Drug (APZ) | Polymer (TAX) | Cross-linker (BaCl<sub>2</sub>) | E.E (%) |
| --- | --- | --- | --- | --- |
| F1 | 10 mg | 10 mg | 5 % | 68.4 % ± 0.65 |
| F2 | 10 mg | 20 mg | 5 % | 75.2 % ± 0.43 |
| F3 | 10 mg | 30 mg | 5 % | 81.5 % ± 1.62 |
| F4 | 10 mg | 40 mg | 5 % | 97.1 % ± 2.36 |
| F5 | 10 mg | 20 mg | 2 % | 51.2 % ± 0.48 |
| F6 | 10 mg | 20 mg | 3 % | 56.8 % ± 0.37 |
| F7 | 10 mg | 20 mg | 4 % | 61.5 % ± 1.45 |

Table 2 reveals an increase in % encapsulation efficiency with the rise in TAX concentration in APZ-TAX-based nanoparticles (F1 to F4). This observed improvement in % encapsulation efficiency can be attributed to the higher viscosity of the solution at increased polymer concentrations, leading to reduced drug diffusion within the polymer matrix [22]. Similarly, for APZ-TAX-based nanoparticles (F5 to F7), lower concentrations of the cross-linker resulted in lower EE. Increasing the cross-linker concentration, however, enhanced the EE. This observation suggests that at lower cross-linker concentrations, the polymer matrix formed may be loose, featuring larger pores due to insufficient cross-linking. This condition leads to higher leaching of drug particles into the solution, resulting in lower EE. Conversely, higher concentrations of the cross-linker, as seen in F4, created a tight network within the matrix and a highly viscous solution, thereby reducing drug leakage and enhancing EE [66]. Based on these findings, F4 emerged as the most optimized formulation among the studied formulations, demonstrating the highest EE.

## Characterization of Optimized TAX based nanoparticles

### Particle Size, Zeta Potential and Polydispersity Index

Within the seven formulations of APZ-TAX-based nanoparticles, the formulation exhibiting the most favourable properties in terms of encapsulation efficiency was identified. The mean particle size, zeta potential, and mean polydispersity index of the optimized formulation (F4) of APZ-TAX-based nanoparticles were assessed.

For poorly soluble drugs, particle size analysis is crucial as it plays a pivotal role in solubility, dissolution, and bioavailability, particularly for APZ-TAX-based nanoparticles. A particle size below 300 nm is essential for effective intestinal transportation into the lymphatic system. The Zeta sizer was employed to analyse particle size, revealing the particle size distribution of the optimized formulation in terms of % intensity, as depicted in Figure 4 (A). The particle size of APZ-TAX-based nanoparticles was determined to be 211.1 nm.

**Figure 4.**
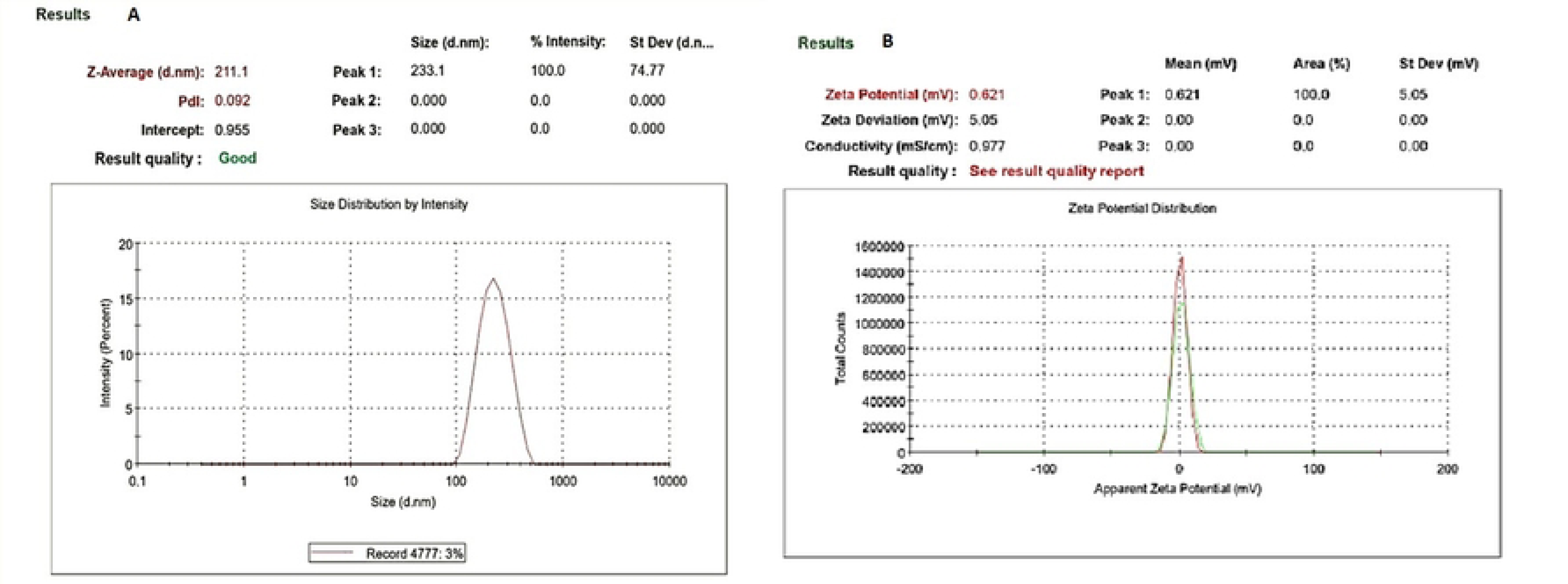
(A) Size distribution of F4 formulation (B) Zeta potential of F4 formulation

A critical indicator for assessing the uniformity of particle size distribution is the Polydispersity Index (PDI). A PDI value below 0.3 indicates a narrow particle distribution, highlighting a homogeneous distribution of nanoparticles. Furthermore, APZ-TAX-based nanoparticles displayed a uniform size distribution, supported by a low PDI of 0.092 and a unimodal size distribution curve, as depicted in Figure 4 (A).

The determination of surface charge and stability of the optimized APZ-TAX-based nanoparticles was conducted by measuring the zeta potential, as illustrated in Figure 4 (B). The recorded zeta potential value for APZ-TAX-based nanoparticles was +0.621 mV, indicating a positive surface charge. This positive zeta potential value (0.621 mV) imparts mucoadhesive characteristics to the TAX-based nanoparticles, attributed to electrostatic interactions between the positively charged nanoparticles and the negatively charged mucin on the mucosal layer. This electrostatic interaction contributes to increased permeability and bioavailability of aripiprazole from TAX-based nanoparticles compared to neutral or negatively charged particles [26]

### Fourier Transforms Infrared Analysis (FTIR)

FTIR analysis was conducted to examine the interactions among APZ, TAX, and the cross-linker. Figure 5 presents the FTIR spectra of TAX, APZ, and the optimized APZ-TAX-based nanoparticle formulation (F4) within the frequency range of 4000–400 cm^-1. The IR spectrum of TAX revealed significant absorption bands, confirming the ester linkage between arabinoxylan and thiourea. Notably, the C=O ester and –SH stretch functional groups were observed at 1649.14 cm^-1 and 2493.96 cm^-1, respectively. The C–O–C stretch appeared at 1170.79 cm^-1, and the stretch at 1031 cm^-1 was noted, with the peak at 821.68 possibly attributed to polymer bending. All FTIR bands were consistent with literature findings.

**Figure 5.**
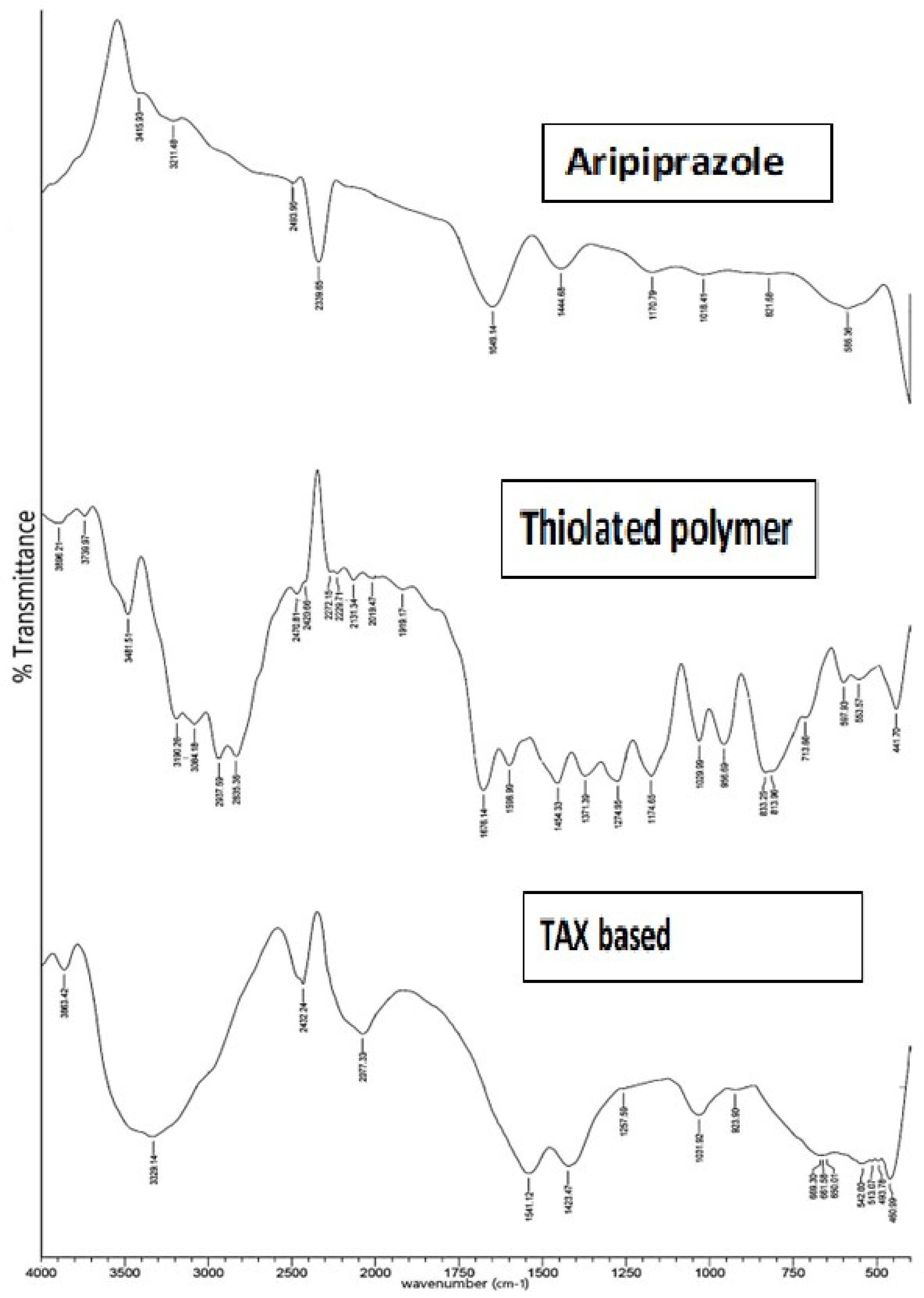
FTIR (A) TAX, (B) APZ, (C) F4 formulation of Aripiprazole TAX based nanoparticles

The FTIR spectrum of pure APZ exhibited peaks representing various functional groups. Absorption bands at 3190.26 and 3481.51 indicated the N-H stretch region, while the peak at 3084.18 represented the C-H aromatic region. Other peaks included C-H aliphatic stretching at 2937.59, C=N (nitrile group) at 2229.71 and 2272.15, C-H bond in aromatic compound at 1676.14, aromatic C-O vibration at 1274.95, and C-Cl stretch region at 553.57, 597.93, 713.66, 813.96, and 833.25. The C-O stretching band was observed at 1174.95.

FTIR analysis was employed to assess the interaction among components in TAX-based nanoparticles. Some characteristic peaks of APZ were absent in the prepared nanoparticles, possibly due to drug entrapment into the matrix. Shifts in certain peaks, such as the N-H stretch from 3481.51 to 3329.14 and the amide group from 1676.14 to 1541.12, indicated hydrogen bond formation between APZ and TAX, confirming their binding. Similarly, peaks at 1274.95, 553.57, and 597.93 experienced slight frequency shifts.

In the case of TAX, some peaks were retained after entrapment, while others shifted slightly. The absorption band at 1649.14 shifted to 1541.12, the absorption band at 1170.79 to 1257, and the absorption band at 821.68 to a slightly higher frequency of 923.9. The FTIR spectral analysis suggests that no strong bonds were formed between the constituents of APZ-TAX-based nanoparticles. However, the presence of hydrogen bonds between APZ and TAX is considered beneficial for establishing interactions within the prepared nanoparticles.

### Differential Scanning Calorimetry (DSC)

The molecular state of APZ in the optimized APZ-TAX-based nanoparticle formulation (F4) was assessed through DSC analysis, as depicted in Figure 6A. DSC thermograms of APZ, thiolated arabinoxylan, and thiolated arabinoxylan-based nanoparticles are presented in Figure 6. The DSC thermogram of APZ exhibited a single, sharp endothermic peak at 140.69 °C, indicative of free crystalline anhydrous APZ. However, in the DSC thermogram of TAX-based nanoparticles, the absence of the endothermic peak of APZ suggests that APZ is fully entrapped within the polymer matrix and has undergone a transformation from its crystalline to amorphous form [27]. The lack of the APZ peak indicates its complete entrapment within the TAX-based nanoparticle formulation, signifying effective absorption in the polymeric network of TAX-based nanoparticles.

**Figure 6.**
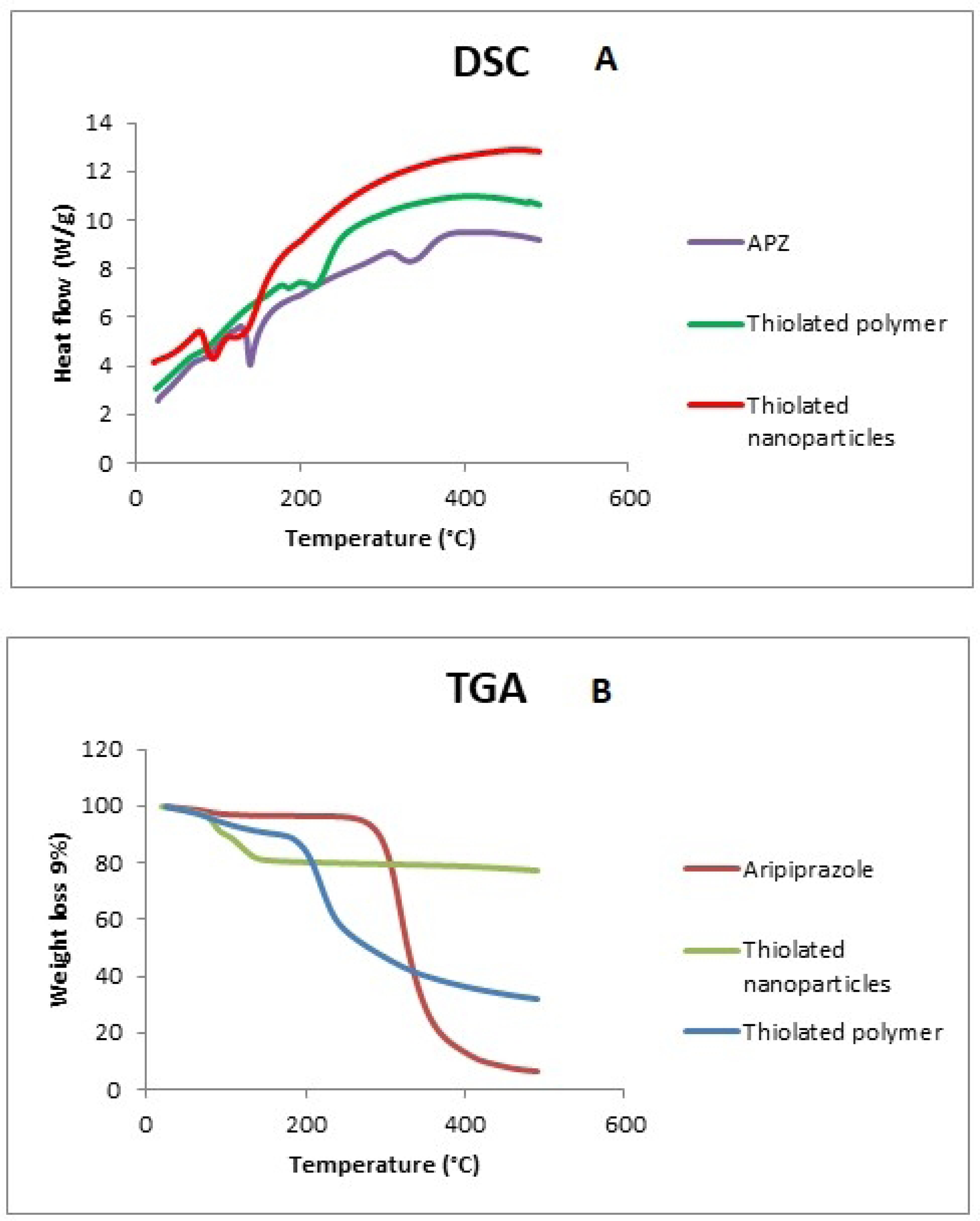
(A) DSC thermogram of pure Aripiprazole, Thiolated polymer, and Thiolated nanoparticle formulation (F4) (B) TGA pattern Aripiprazole, Thiolated polymer, and Thiolated nanoparticle formulation (F4)

The DSC thermogram of TAX showed a peak at 216.24 °C, corresponding to its melting point. The disappearance of this peak in the DSC thermogram of thiolated arabinoxylan-based nanoparticles revealed the complete transformation of thiolated arabinoxylan into the amorphous form. These findings provide evidence that APZ underwent a transformation to an amorphous state during the loading procedure [28]. It’s noteworthy that the conversion of crystalline APZ into an amorphous form within TAX-based nanoparticles contributes to improved aqueous solubility and bioavailability [29].

### Thermogravimetric analysis (TGA)

The TGA curves for pure APZ, TAX, and the optimized formulation (F4) of TAX nanoparticles are illustrated in Figure 8. TGA is employed to quantify the total concentration of APZ entrapped in the polymer matrix. In this study, APZ desorption after decomposition is represented as temperature-dependent weight loss [30].

**Figure 7.**
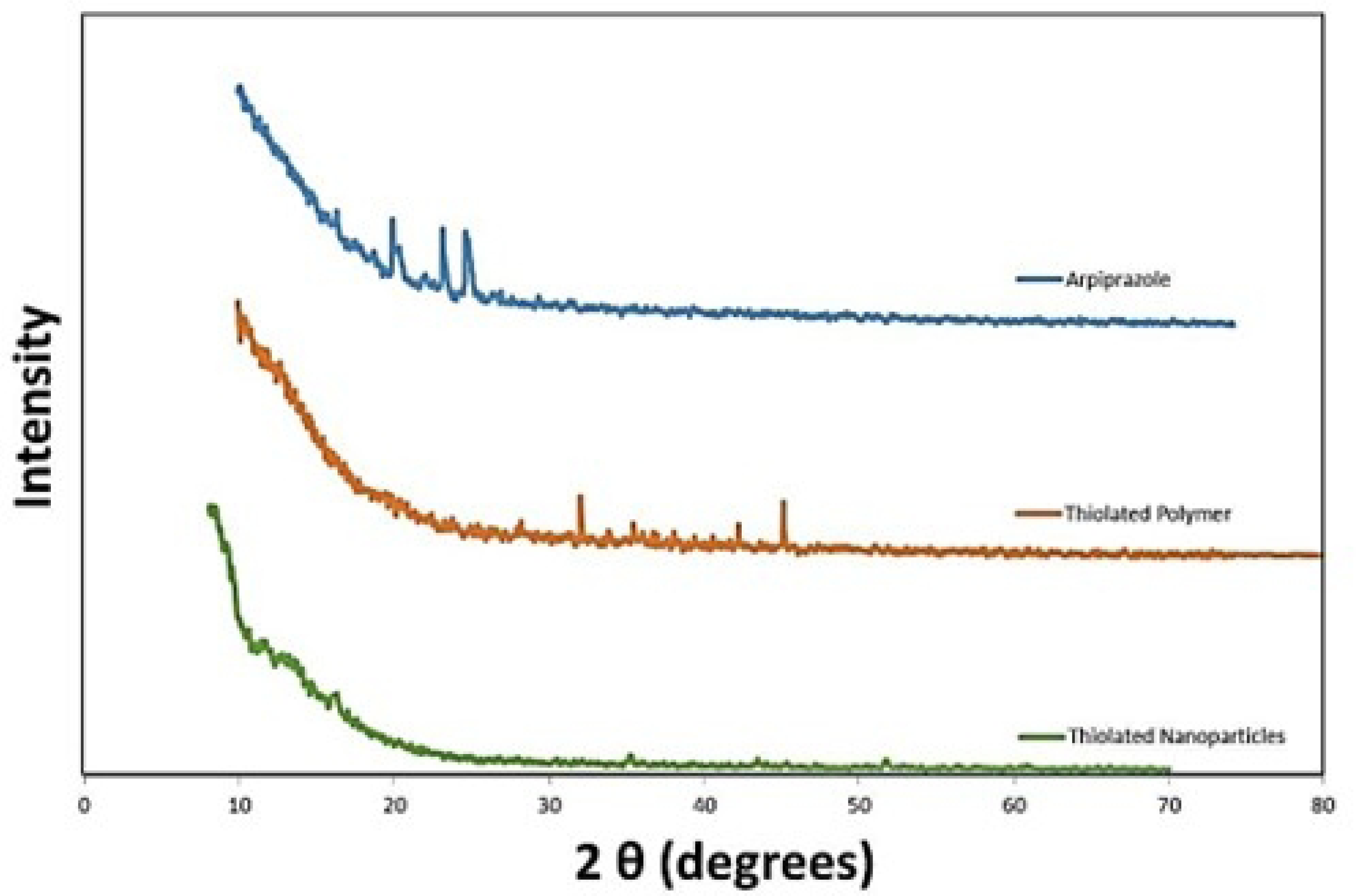
Powder X-ray diffraction patterns of Aripiprazole, Thiolated polymer (arabinoxylan) and Thiolated nanoparticle formulation (F4)

**Figure 8.**
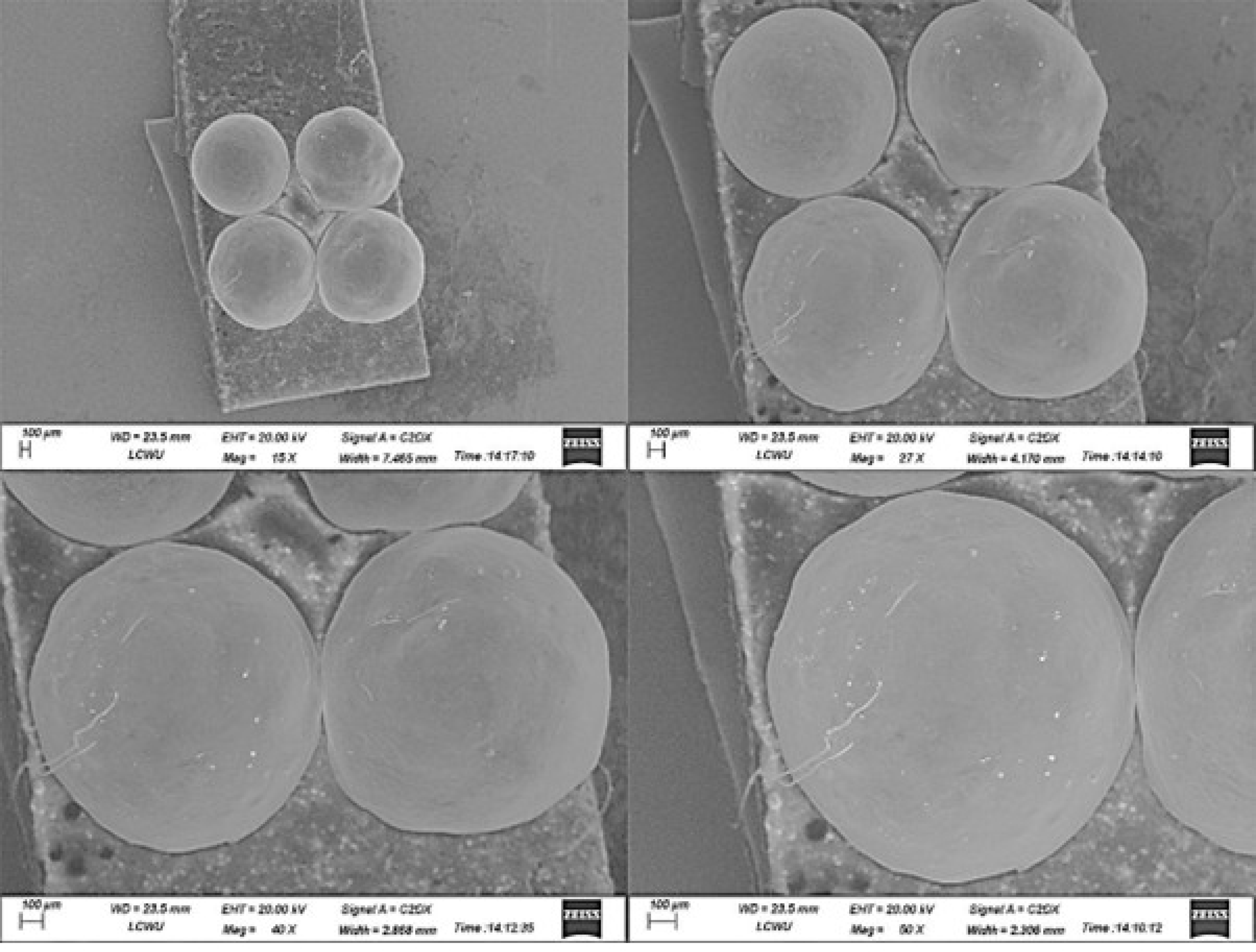
Scanning electron microscopy images of F4 formulation of Thiolated arabinoxylan based nanoparticles

The estimation of APZ loaded concentration in TAX-based nanoparticles was determined by the ratio of weight loss to the initial weight between 100 °C to 95.89 °C. A gradual weight loss was observed with an increase in temperature, and APZ degradation occurred in three stages as shown in if figure 6B. Initially, there was a 4.11% weight loss at 95.89 °C, followed by a second stage involving a more substantial reduction in weight of 77.81% at 376.14 °C. The last stage of decomposition occurred at 474.12 °C, accounting for 11.151% weight loss. Similarly, the weight loss of TAX also occurred in three steps. The initial decomposition took place at 182.07 °C, indicating a weight loss of 11.09%. The second stage involved a weight loss of 29.35% at 239.39 °C. The final stage of decomposition occurred at 465.1°C, attributed to TAX-based nanoparticles, confirming the entrapment of APZ within TAX-based nanoparticles.

### Powder X-ray Diffractogram (PXRD)

The physicochemical characteristics of the entrapped drugs were assessed through PXRD. Figure 7 illustrates the physical states of APZ, TAX, and APZ-TAX-based nanoparticle formulation (F4). In the PXRD study of the optimized APZ-loaded TAX-based nanoparticle formulation (F4), the XRD diffractogram containing various sharp peaks of the pure drug indicated the highly crystalline form of the drug. The disappearance or significant reduction in the sharp drug peaks signifies the amorphous nature of the drug [31].

PXRD analysis was employed to characterize the crystalline and amorphous structural assembly within APZ, TAX, and thiolated arabinoxylan-based nanoparticles. The PXRD analysis revealed that APZ lost its crystalline form upon being entrapped within the TAX-based nanoparticles. The X-ray diffractogram of APZ depicted sharp peaks at different scanning 2θ degrees, including 19.96°, 23.2°, and 24.68°, along with various secondary peaks representing its crystalline form [32]. Similarly, TAX exhibited sharp peaks at 32.12° and 45.2°, as well as a few diffused peaks similar findings were observed in a previous study where they explored the pharmaceutical application of TAX, [33]. In contrast, PXRD of thiolated arabinoxylan-based nanoparticles did not exhibit intense peaks of APZ and TAX suggesting a decrease in crystallinity and presence of their amorphous form. This correlates with the enhanced aqueous solubility of APZ from TAX-based nanoparticles [34].

### Scanning Electron Microscopy (SEM)

The surface and cross-sectional micrographs of the optimized APZ-TAX-based nanoparticle formulation (F4) are displayed in Figure 8. Scanning electron microscopy (SEM) was employed to evaluate the morphology of TAX-based nanoparticles at various magnifications. From the SEM images, it was observed that the prepared TAX-based nanoparticles were discreetly spherical with a uniform morphology. The SEM analysis revealed that the uniform surfaces exhibited small bright spots. These bright spots on the grey background corresponded to pores on the surfaces of TAX-based nanoparticles. The porous surface of TAX-based nanoparticles is a typical shape of polymeric particles formed during the drying of droplets. This could be attributed to solvent evaporation, leading to the shrinkage of the polymeric layer on the droplet surface [35]. The presence of pores on the surfaces of TAX-based nanoparticles serves as solvent entry channels, facilitating the release of the drug from these nanoparticles upon contact with an aqueous medium [36].

### In-Vitro Dissolution and Drug Release Kinetics

An in-vitro release study of aripiprazole (APZ) from all seven APZ-TAX-based nanoparticle formulations was conducted in both simulated gastric fluid (SGF) and simulated intestinal fluid (SIF) at 37 °C ± 0.5 °C using the USP type II dissolution apparatus with stirring at 50 rpm. Thiolated arabinoxylan-based nanoparticles exhibited pH-dependent release characteristics, and the results were graphically presented in Figure 9. The release study of APZ from all the formulations was studied in both dissolution media [37].

**Figure 9.**
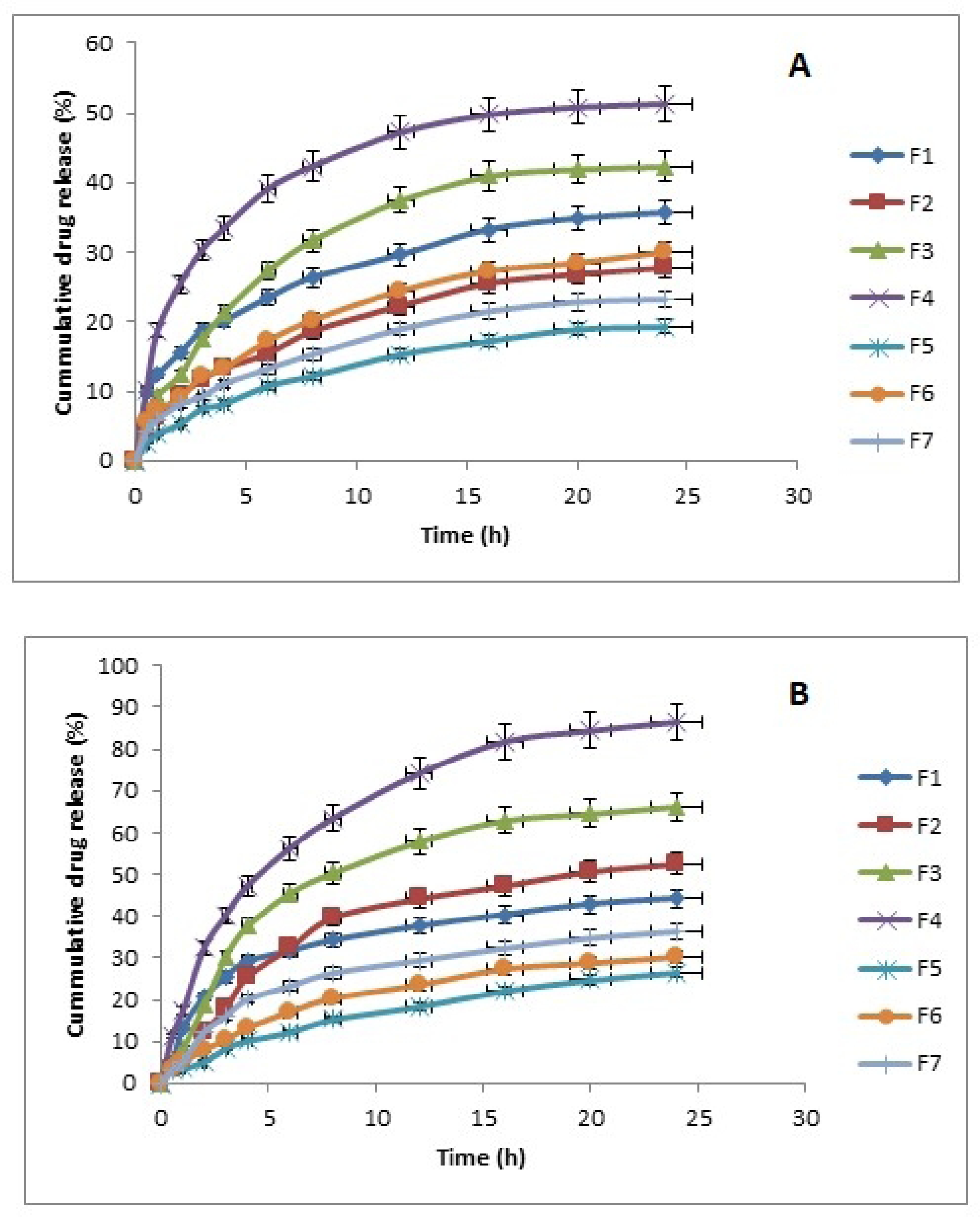
In vitro drug release profile of thiolated arabinoxylan based nanoparticles (F1 to F7) (A) SGF buffer (B) SIF buffer

In SGF, TAX-based nanoparticles F1, F2, F3, F4, F5, F6, and F7 exhibited releases of 51.27%, 42.13%, 55.16%, 64.32%, 26.24%, 36.44%, and 31.89%, respectively. In the first 30 minutes, F3 showed a burst release of drug of about 32.41%, followed by sustained release of about 55.16%, and F4 showed a burst release of about 46.44%, followed by sustained drug release of about 64.32%. However, in SIF, APZ-loaded TAX-based nanoparticle formulations (F1, F2, F3, F4, F5, F6, and F7) showed maximum releases of 51.68%, 55.45%, 69.21%, 79.22%, 25.53%, 31.82%, and 41.56%, respectively, after 24 hours. At this pH, the rate of drug release was significantly higher, and the drug was released in a sustained manner. The slow release of APZ after some time was attributed to the swelling of the polymer or its slow degradation. Some part of APZ in TAX-based nanoparticles was not released, possibly because the particles were not fully dissolved or disintegrated in the dissolution medium. This feature can be advantageous as it prevents the drug from entirely degrading in the stomach and promotes the intestinal absorption of APZ. Hence, more undegraded drug moves towards the small intestine [38]. TAX-based nanoparticles were stable in the intestinal environment and exhibited prolonged intestinal residence time, facilitating drug absorption from the intestine [39].

A high concentration of cross-linker would produce a high degree of cross-linking, inhibiting the penetration of aripiprazole into the polymer matrix and leading to the desorption of aripiprazole from the particle surface, thereby increasing its release. A low degree of cross-linking would cause increased diffusion of aripiprazole from the particle surface toward its core matrix compared to more tightly cross-linked samples [40].

The in-vitro drug release kinetics of APZ-loaded TAX-based nanoparticles (F1-F7) were systematically investigated to identify the most suitable release behaviour as indicated in Table 3. Comparing the regression coefficient (R2) values in both scenarios enabled the identification of the most suitable kinetic model for drug release. A correlation coefficient value (R2) close to 1 indicates a well-fitted model for the drug release mechanism [41]. In the case of simulated gastric fluid (SGF), all APZ-loaded TAX-based nanoparticle formulations (F1-F7) were found to follow the Korsmeyer-Peppas model. The kinetic data obtained suggested that the n value (diffusion coefficient) for all TAX-based nanoparticle formulations was equal to or less than 0.45, indicative of Fickian diffusion. Notably, F5 TAX-based nanoparticle formulation exhibited an n value greater than 0.45, indicating non-Fickian diffusion. In the context of simulated intestinal fluid (SIF), the kinetic data revealed that all APZ-loaded TAX-based nanoparticle formulations followed the Korsmeyer-Peppas model as the best-fit model for drug release. Specifically, the F6 formulation adhered to both the Higuchi and Korsmeyer-Peppas models. Further analysis of the kinetic data indicated that the n values obtained were greater than 0.45 for F2, F3, F5, and F6 TAX-based nanoparticle formulations, suggesting non-Fickian diffusion. In contrast, F1, F4, and F7 formulations followed non-Fickian diffusion, as evidenced by n values less than 0.45.

**Table 3.**
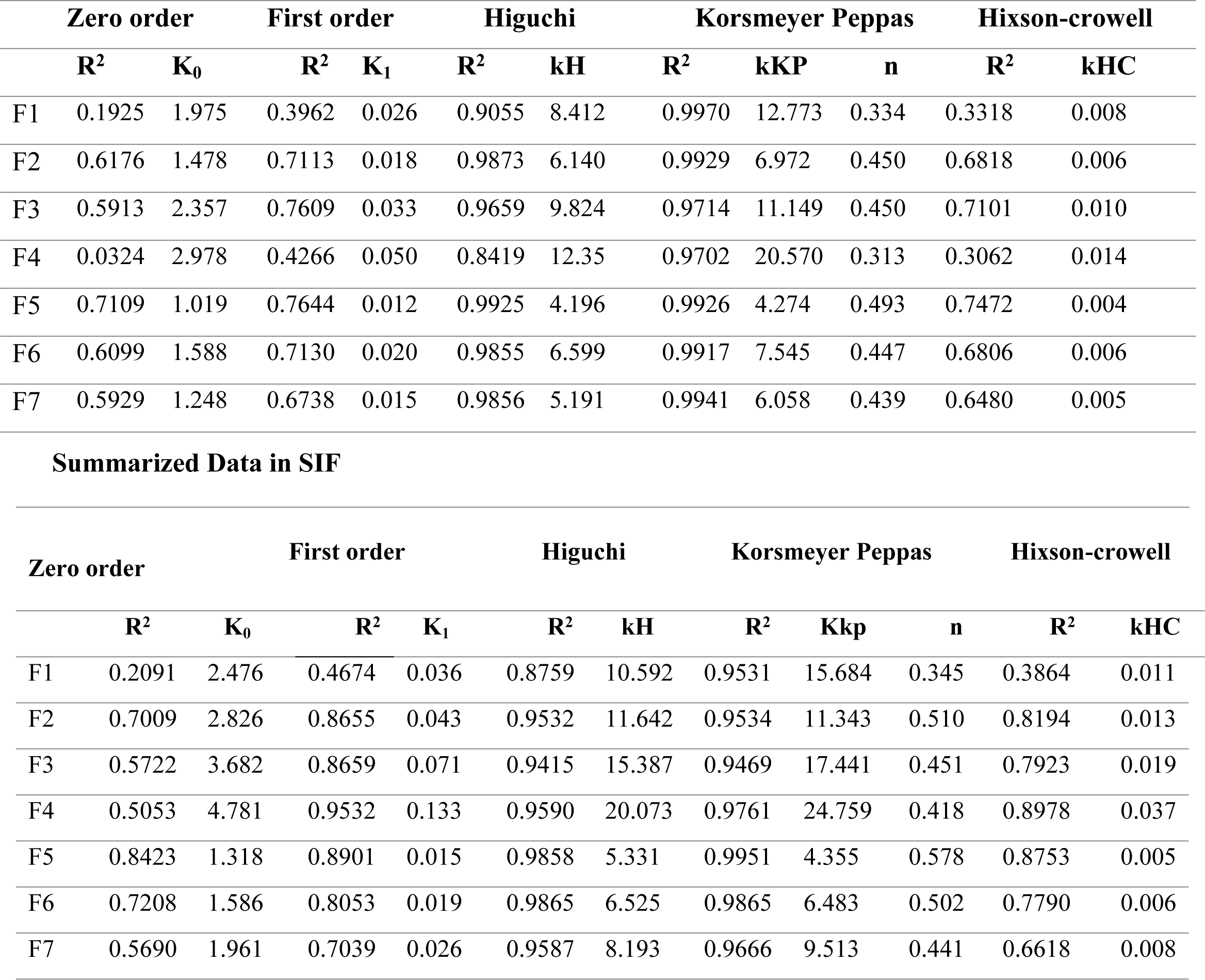
Correlation Coefficients (R^2^) and Release Rate Constants of Thiolated arabinoxylan based nanoparticles in SGF and SIF.

### Ex-vivo intestinal Permeation Study

Ex vivo permeation studies are conducted to investigate the transport of drugs or formulations across intestinal membranes using excised tissues. In this context, the everted sac method is utilized to assess the intestinal permeation profiles of APZ-loaded TAX-based nanoparticle formulations (F1-F7) in simulated gastric fluid (SGF) and simulated intestinal fluid (SIF). Figure 10 illustrates the intestinal permeation profiles of APZ-loaded TAX-based nanoparticle formulations (F1-F7) in both simulated gastric fluid (SGF) and simulated intestinal fluid (SIF) using the everted sac method. In SGF, depicted in Figure 10 (A), the cumulative amount of aripiprazole from F1-F7 formulations permeating the intestinal wall to the buffer after 120 minutes was observed as follows: 64.432%, 49.442%, 71.023%, 79.223%, 33.884%, 49.442%, and 40.646%, respectively.

**Figure 10.**
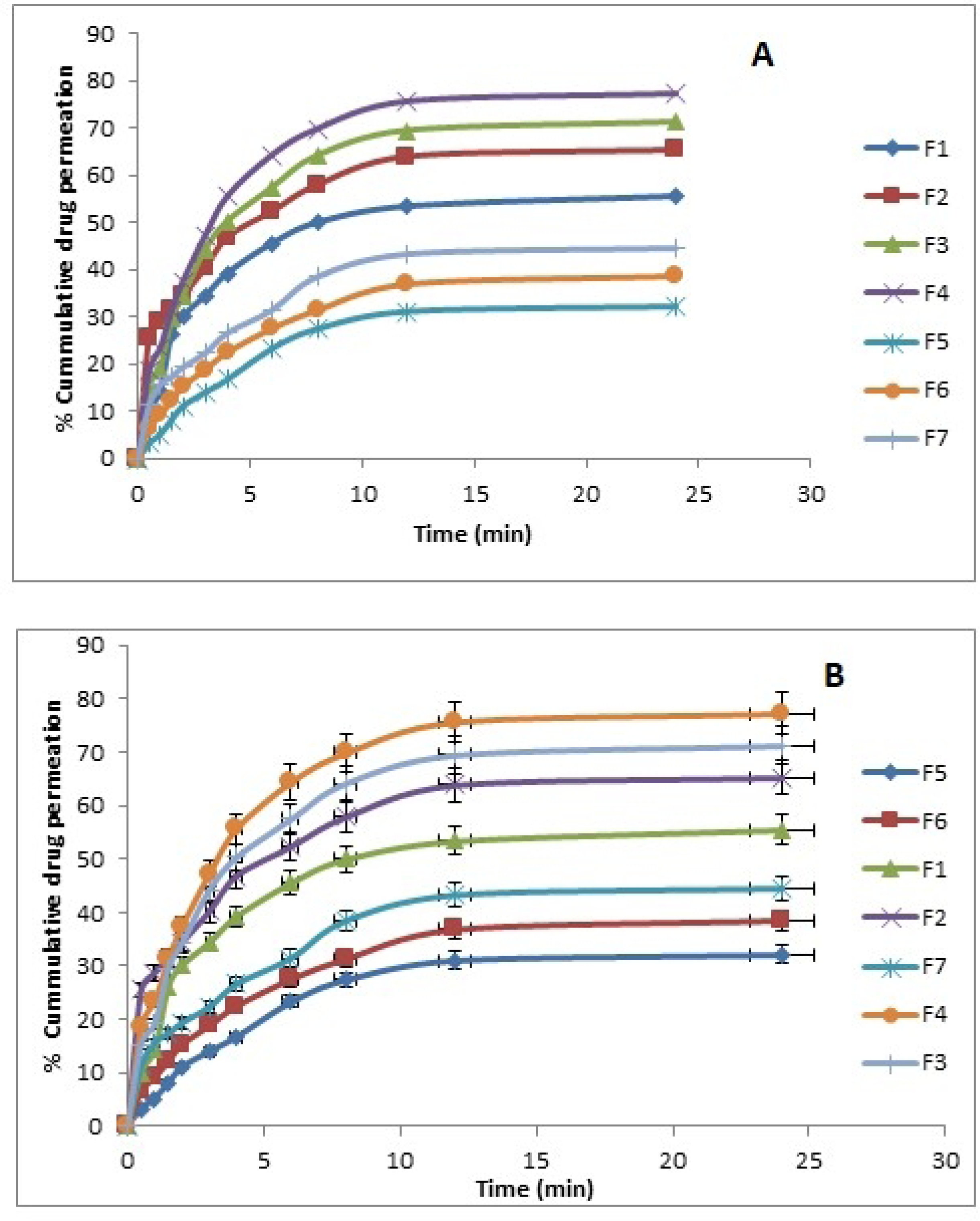
Ex vivo gut-permeation profile showing comparative cumulative drug transport (A) SGF (B) SIF

In the case of SIF, represented in Figure 10 (B), the cumulative amount of aripiprazole from APZ-loaded TAX-based nanoparticle formulations F1-F7 permeating through the intestine to the buffer after 120 minutes was found to be 55.477%, 65.244%, 71.226%, 77.234%, 32.211%, 38.4355, and 44.536%. The figures indicate that, in both SGF and SIF cases, a greater amount of APZ was delivered from the F4 TAX-based nanoparticle formulation through the intestinal mucosa compared to others. This substantial increase in absorption through the intestinal mucosa may be attributed to the increased dissolution of APZ from the F4 formulation after incorporation into the matrix of TAX-based nanoparticles. Additionally, the nanosized formulation contributes to enhanced permeation through the intestinal wall. Moreover, the inhibition of P-glycoprotein (P-gp) expression in the gut lumen by P-gp inhibitors, such as TAX, enhances drug absorption, ultimately increasing the drug’s bioavailability [42].

### In-vivo Pharmacodynamic Study

Figure 11 indicates pictorial representation of forced swim test, tail suspension test, and elevated plus maze test whereas figure 12 indicates graphical representation of catalepsy test, passive avoidance test, forced swim test, and tail suspension test.

**Figure 11.**
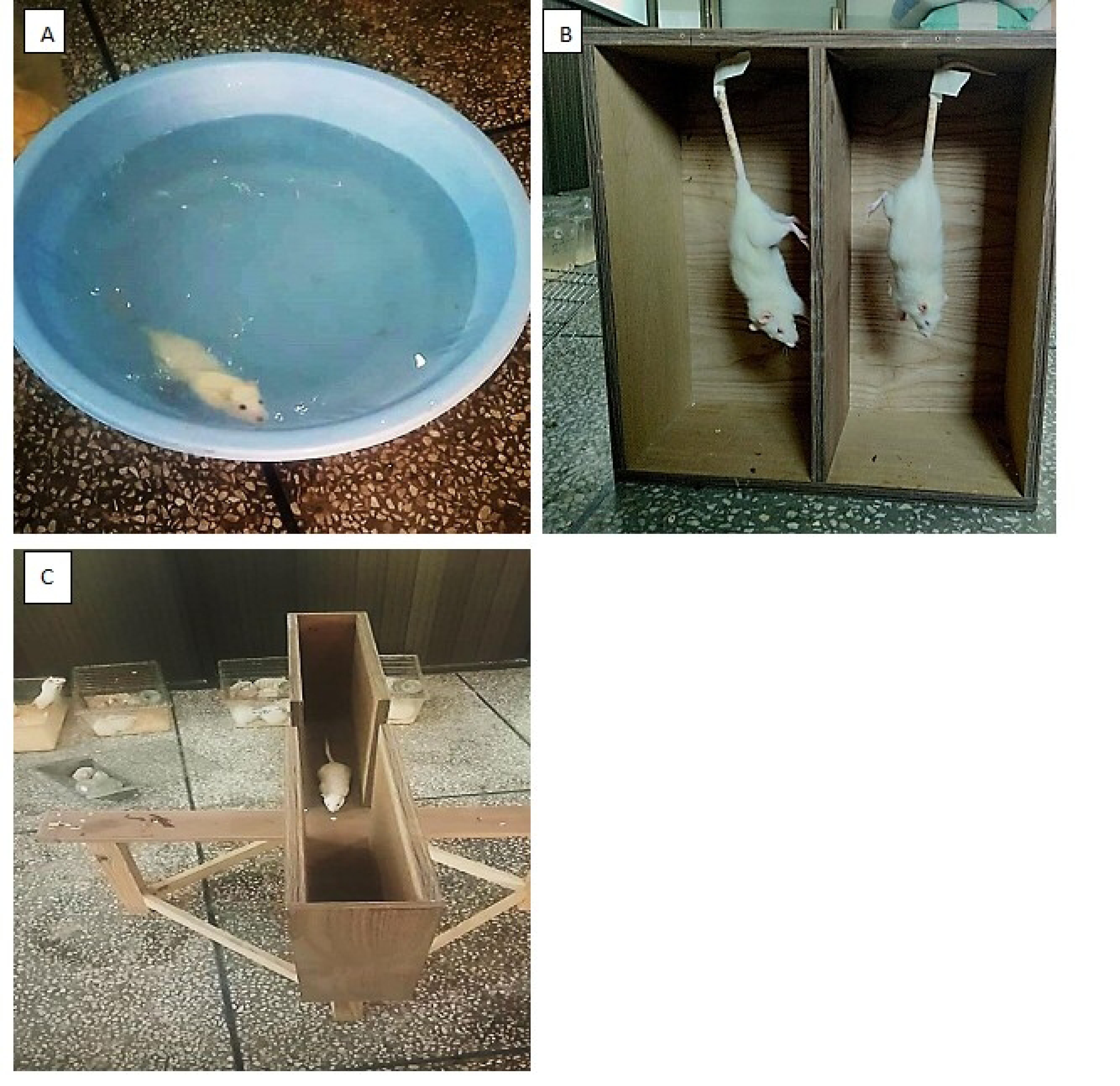
Pictorial representation of (A) Forced swim test (B) Tail suspension test and (C) Elevated pulse maize test

**Figure 12.**
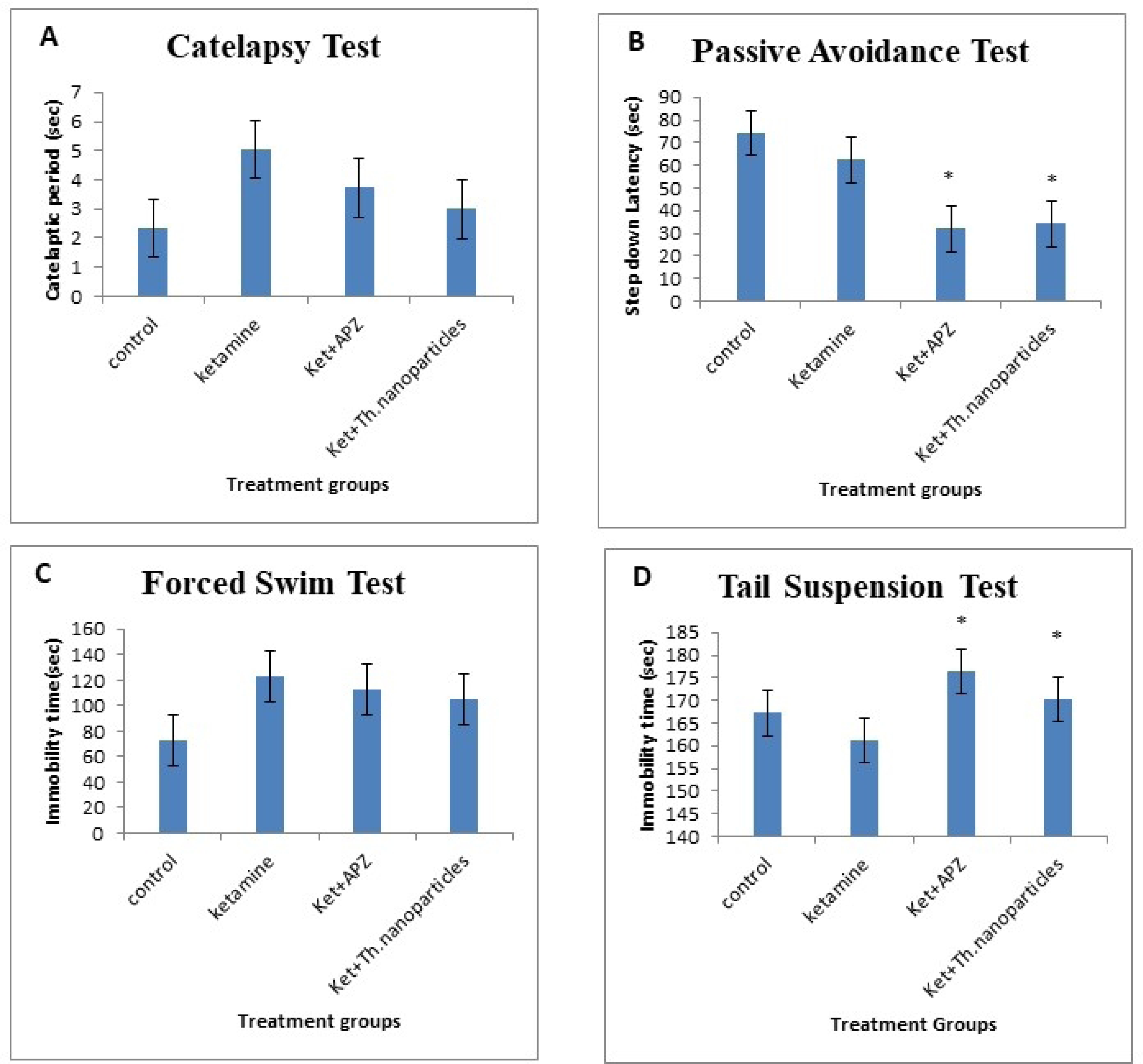
Effects of Ketamine alone, Ket+APZ and Ket+TAX based nanoparticles on the behavioural performance of rats. Ketamine was administered i.p., 30 min before the session. Values are mean ± S.E.M. Data are presented graphically (A) Catelapsy test (B) Passive avoidance test (C) Forced swim test (D) Tail suspension test. ‘*’ p ≤ 0.05 against control group.

### Catalepsy Test

The catalepsy test assesses the potential motor side effects of drugs, and in the case of aripiprazole, a commonly prescribed antipsychotic, it aims to evaluate its impact on motor skills and the risk of inducing cataleptic responses in experimental animals. The administration of pure oral APZ suspension (APZ 3mg/kg) and oral APZ-TAX-based nanoparticles (APZ 3mg/kg) did not elicit any marked effects on the motor skills of rats. Post-hoc analysis using ANOVA revealed that both the Ket + APZ and Ket + APZ+TAX based nanoparticles groups exhibited no significant differences when compared with the control group, with significance values (P < 0.05). In rats subjected to chronic administration of APZ suspension and the optimized APZ-TAX-based nanoparticle formulation (F4) previously treated with Ketamine (50 mg/kg i.p.), no cataleptic effects were observed, indicating the absence of motor skill impairment compared to the control group [43].

### Passive Avoidance Test

The passive avoidance test gauges memory retention by assessing an animal’s ability to recall and avoid an aversive stimulus, providing insights into the cognitive effects of substances like aripiprazole. The results of passive avoidance test are shown in figure 12 (B). A 14-day administration of Ketamine (50 mg/kg, i.p.) resulted in a reduction in step-down latency, indicating memory impairment. Treatment with pure APZ suspension (APZ 3 mg/kg, p.o.) and TAX-based nanoparticles (APZ 3 mg/kg, p.o.) mitigated the ketamine-induced increase in step-down latency, with both exhibiting statistical significance (P > 0.05) compared to the control group [44].

### Forced Swim Test

The Forced Swim Test on the 15th day evaluated the impact of APZ and its APZ-TAX-based nanoparticle formulation (F4) in a Ketamine-induced schizophrenia model. Chronic intra-peritoneal administration of ketamine for 14 days substantially increased the immobility period compared to the control group (P < 0.001), an effect significantly attenuated by the pure APZ suspension group (APZ 3 mg/kg) administered orally and previously treated with ketamine (P < 0.05) as shown in 12 (C). Similarly, the APZ-TAX-based nanoparticles group (APZ 3 mg/kg) administered orally, previously treated with ketamine at 50 mg/kg, also significantly reduced the immobility period (P < 0.05) [45].

### Tail Suspension Test

T Tail suspension test conducted on the 15th day revealed that prolonged administration of Ketamine (50 mg/kg i.p.) for 14 days significantly decreased immobility time as shown in figure 11 (B). The groups treated with oral APZ suspension (APZ 3 mg/kg) and oral Thiolated arabinoxylan-based nanoparticles (APZ 3 mg/kg), both previously subjected to ketamine (50 mg/kg i.p.), displayed immobility durations almost comparable to the control group, indicating a significant reduction (P < 0.05) as shown in figure 12 (D). Two-way ANOVA demonstrated a substantial impact of Ketamine (P ˃ 0.05). Similarly, significant effects were also observed in the APZ groups previously treated with ketamine (P < 0.05) and APX-TAX-based nanoparticle formulation (F4) previously treated with ketamine (P < 0.05) [46].

### Elevated Pulse Maze Test

The Elevated Plus Maze Test, conducted on the 15th day, assessed the effects of Ket+APZ Suspension (APZ 3 mg/kg p.o) and Ket+APZ TAX-based nanoparticles (APZ 3 mg/kg p.o). Both groups, exhibiting significance at P < 0.05, demonstrated significant efficacy in improving animal behaviour, manifesting moderate anxiolytic effects. These effects were characterized by an increased time spent in open arms and a decreased time spent in closed arms, as illustrated in Figure 13 (A) and Figure 13(B). Notably, both groups exhibited a tendency to increase the number of open arm entries compared to the control (P < 0.05), as depicted in Figure 13(C) [47].

**Figure 13.**
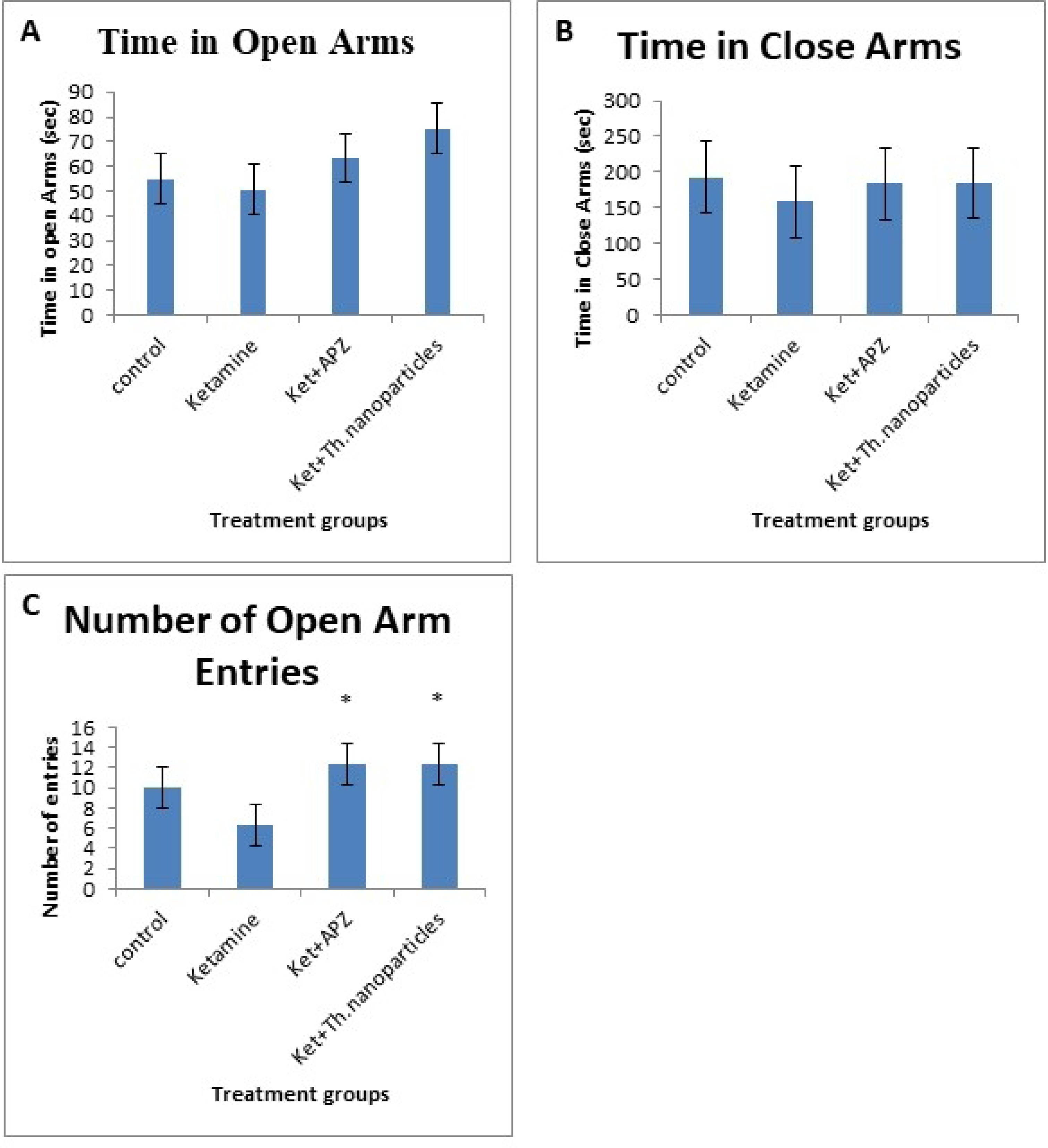
Elevated Plus Maze Test: Anxiety Measurement in Rats Treated with Ketamine Alone, Ket+APZ, and Ket+Thiolated Nanoparticles A) Number of Open Arm Entries; B) Time Spent in the Close Arms; C) Time Spent in the Open Arms.

### Acute Oral Toxicity Study

The in-vivo acute oral toxicity study of the APZ-loaded TAX-based nanoparticle formulation (F4) spanned 14 days in male rats, administering an oral dose of 5000 mg/kg body weight. Throughout the study, both the control and treatment group rats exhibited insignificant changes in weight gain after the oral administration of 5000 mg/kg TAX-based nanoparticles and APZ. Table 4 documents the effects of TAX-based nanoparticles and aripiprazole intake, along with nutrient intake, on body weights. No mortality or signs of illness, such as a runny nose, eye irritation, or vomiting, were observed in any rat from either the control or treatment group. Similarly, none of the rats showed signs of acute oral toxicity, and all vital organs, including the heart, kidneys, lungs, liver, brain, stomach, and spleen, remained unaffected. Physical parameters, including faeces consistency, eyes, fur and skin condition, urine colour, sleep, and salivation, remained normal throughout the study. According to the Globally Harmonized System (GHS) for the testing of chemicals, the LD50 value exceeding 2g/kg dose falls under category 5 with a toxicity score of zero. Consequently, TAX-based nanoparticles align with category 5, indicating zero toxicity. The acute toxicity study aimed to evaluate potential toxic effects from arabinoxylan and TAX. The results demonstrated that rats exhibited no signs of toxicity. Over the 14-day period, no mortality occurred in any rat from the five groups, and both treated and non-treated groups displayed no toxic effects. Physical observations indicated no alterations in behavioural patterns, fur, sleep patterns, or eyes of the rats. Body weights were recorded before dosing, on the 3rd, 7th, and final day, revealing a gradual increase in the body weight of the treated group until the 14th day, with slight weight gain observed in both control and treated groups [48].

**Table 4.** Clinical Observations for Acute Oral Toxicity testing of Group 1, Group 2, Group 3, and Group 4.

| Observations | Group 1<br>(Control) | Group 2<br>(Ketamine) | Group 3<br>(Ket+APZ<br>Suspension) | Group 4<br>(Ket+TAX based<br>nanoparticles) |
| --- | --- | --- | --- | --- |
| Signs of illness | Nil | Nil | Nil | Nil |
| Body Weight (g) |  |  |  |  |
| Day-1 | 173.33±7.32 | 168.36±4.43 | 187±8.26 | 191±7.46 |
| Day-3 | 177.12±8.91 | 170.84±2.22 | 191.64±2.98 | 201.33±8.98 |
| Day-7 | 181.66±4.64 | 174.36±5.11 | 199.37±3.41 | 211.54±2.96 |
| Day-14 | 187.35±2.14 | 182.86±8.91 | 210.64±1.65 | 224.32±6.31 |
| Food consumption (g) |  |  |  |  |
| Day-1 | 86.00±3.26 | 87.32 | 84.06±2.68 | 89.33±6.12 |
| Day-3 | 91.34±2.34 | 90.32±2.45 | 81.65±3.26 | 90.08±8.33 |
| Day7 | 75.23±8.81 | 78.34±2.14 | 74.52±8.23 | 78.21±7.26 |
| Day-14 | 64.53±6.35 | 69.34±5.37 | 70.00±3.28 | 66.38±3.91 |
| Water intake (mL) |  |  |  |  |
| Day-1 | 228.13±1.86 | 238.27± 7.91 | 232.26±12.37 | 235.22±2.48 |
| Day-3 | 234.35±6.62 | 237.66±7.13 | 241.44±4.62 | 234.66±5.23 |
| Day-7 | 210.23±5.42 | 214.61±9.81 | 220.34±3.55 | 212.25±4.13 |
| Day-14 | 196.21±1.92 | 190.33±4.46 | 202.33±3.43 | 196.55±4.67 |

On the 15th day of the experimental study, all rats from each group were sacrificed for a comprehensive organ assessment. The seven vital organs—heart, brain, liver, stomach, spleen, kidney, and lung—underwent examination to detect any abnormalities. No significant alterations were observed, and the absence of inflammation or lesions in the histopathological assessment of the heart, brain, liver, kidney, lung, and spleen strongly indicated the safety of APZ and TAX [49].

### Histopathological Examination

All major organs, encompassing the heart, brain, kidney, liver, stomach, lung, and intestine, underwent a thorough washing with normal saline and were subsequently preserved in 10% buffered formalin for histopathological evaluation across all four experimental groups. Subsequent immersion in paraffin enabled the staining of crucial rat tissues with haematoxylin and eosin, facilitating detailed histopathological analysis. Capturing images at a magnification of ×400 for both treated and non-treated groups, the observations revealed the absence of toxic effects in vital organs for both AX and TAX as indicated in Figure 14.

**Figure 14.**
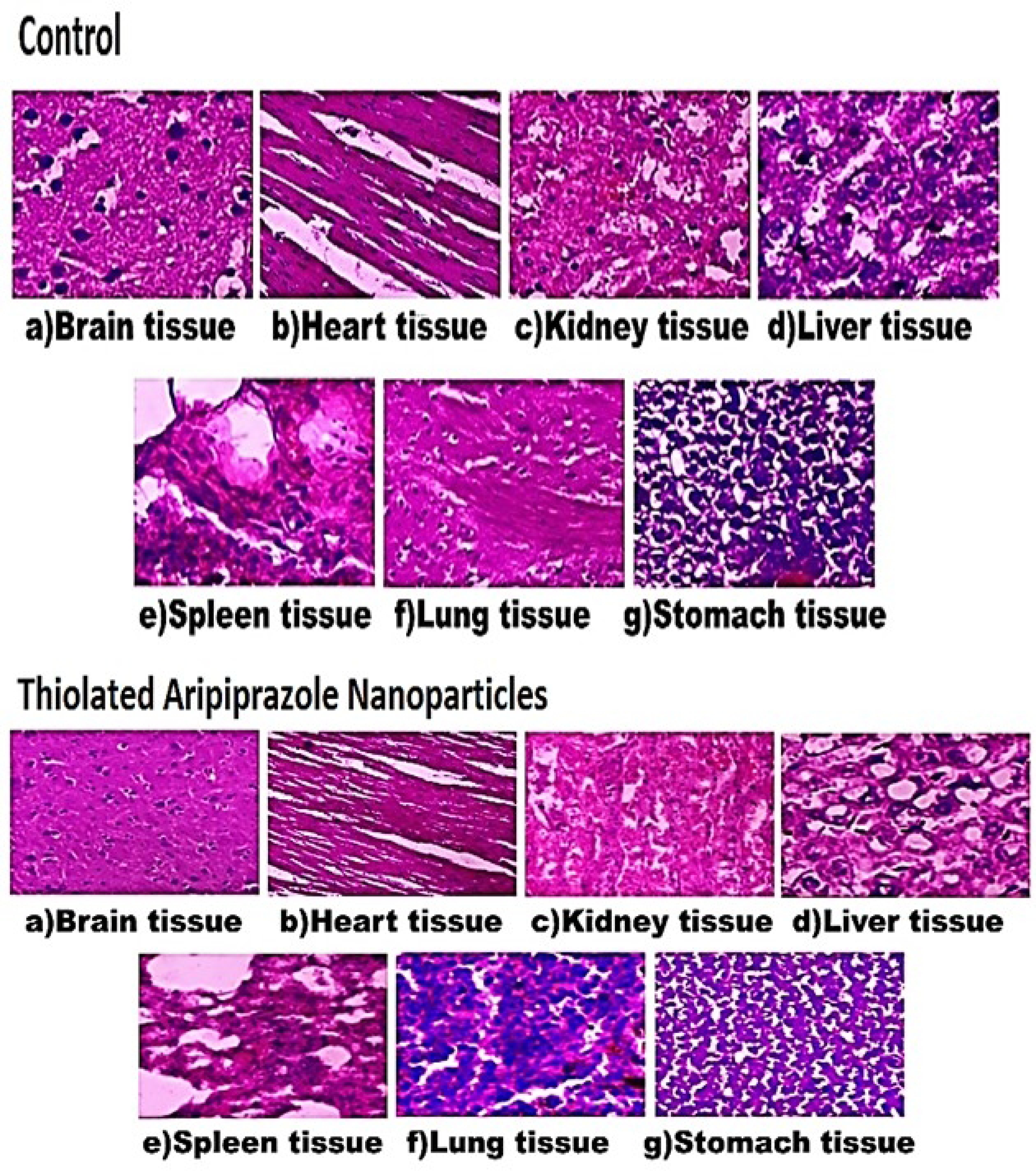
Comparative histopathological analysis of tissues from vital organs in an acute oral toxicity study: control group vs. Thiolated aripiprazole nanoparticle formulation group

Specifically, the myocardium tissues exhibited integrity, and the liver displayed a normal lobular architecture, with mild inflammation observed in portal tracts. Standard pathology assessment of stomach mucosa, spleen, brain, lung, and kidney disclosed no signs of deterioration. Consequently, all vital organs exhibited exemption from significant pathology, and clinical findings for both control and treated rats fell within the normal range [50].

The investigation meticulously assessed the effects of ketamine, aripiprazole (APZ) in suspension, and APZ-loaded thiolated arabinoxylan (TAX)-based nanoparticles on a spectrum of haematological and biochemical markers, detailed in Table 6 [51].

**Table 6.** Biochemical Blood Analysis of Group 1, Group 2. Group 3, and Group 4.

| Parameters | Group 1<br>(Control) | Group 2<br>(Ketamine) | Group 3<br>(Ket+APZ<br>Suspension | Group 4<br>(Ket+TAX based<br>nanoparticles) |
| --- | --- | --- | --- | --- |
| Hemoglobin (g/dL) | 12.2±4.6 | 12.1±3.7 | 13.3±4.1 | 14.3±2.5 |
| WBCs (10<sup>3</sup>/mm<sup>3</sup>) | 6.1±1.32 | 5.4±4.21 | 6.2±6.11 | 5.1±0.88 |
| RBCs (mL/mm<sup>3</sup>) | 4.8± 3.75 | 4.8± 3.75 | 7.3± 3.24 | 8.0± 2.46 |
| Platelets (10<sup>3</sup>/mm<sup>3</sup>) | 396±4.1 | 432±3.5 | 543±2.2 | 650±4.3 |
| Hematocrit (%) | 44±2.6 | 45±7.1 | 47±3.4 | 45±4.6 |
| MCV (fL) | 80±3.76 | 79±4.21 | 85±2.31 | 83±3.42 |
| MCH (Pg) | 24±1.54 | 28±0.51 | 15±0.51 | 18±0.42 |
| MCHC (g/dl) | 33±3.21 | 30.1±2.35 | 34±3.44 | 31.6±1.43 |
| ALT (U/L) | 38±1.99 | 33±0.72 | 44±2.11 | 55±0.13 |
| AST (U/L) | 34±2.35 | 41±3.42 | 38±0.24 | 39±0.13 |
| Total Bilirubin (mg/dl) | 0.4 ±4.11 | 0.2±1.32 | 0.7±2.11 | 0.6±3.21 |
| Creatinine (mg/dl) | 0.7±2.65 | 0.6±4.11 | 0.5±3.52 | 0.4±1.24 |
| Urea (mg/dl) | 32±3.22 | 28±4.21 | 34±4.11 | 37±2.46 |
| Uric acid (mg/dl) | 5.6±4.31 | 6.2±3.70 | 5.1±0.81 | 4.6±3.04 |
| Cholesterol (mg/dl) | 162±0.73 | 166±0.88 | 148±3.24 | 136±1.60 |
| Triglyceride (mg/dl) | 121±2.13 | 90±2.80 | 104±3.12 | 86±1.44 |
| HDL (mg/dl) | 48±2.14 | 41±3.34 | 56±3.25 | 63±1.26 |
| LDL (mg/dl) | 132±2.62 | 124±4.62 | 96±3.25 | 90±2.64 |

Haematological scrutiny revealed an upward trend in haemoglobin levels for groups treated with TAX nanoparticles loaded with APZ or APZ solution in the presence of ketamine, suggesting a potential enhancement of oxygen transport. Increased platelets count in groups receiving APZ formulations indicated a plausible modulation of clotting processes. Notably, measurements related to red blood cells (RBC), such as haematocrit and count, exhibited an upward trend in groups administered with APZ formulations, which is insignificant when compared to control.

White Blood Cell (WBC) count, a key indicator of immunological response, exhibited no significant differences across groups, indicating negligible impact on immune function by the formulations. Liver function indicators, including ALT and AST, remained within the normal range, indicating an absence of hepatotoxicity. Levels of total bilirubin remained largely unchanged. Renal function parameters, such as urea and creatinine levels, displayed similarities across groups, signifying that APZ formulations did not induce substantial changes in renal function. Metabolic and lipid indicators, including triglycerides and lipoprotein levels, fell within the normal physiological range, with slight deviations noted. In groups treated with APZ formulations, HDL cholesterol exhibited an upward trend, suggesting a potential positive impact on cardiovascular health.

In summary, this comprehensive evaluation of haematological and biochemical characteristics in the presented table implies that TAX nanoparticles loaded with APZ, particularly in conjunction with ketamine, may influence specific blood and metabolic parameters. Crucially, observed alterations remained within physiological bounds, emphasizing the relative safety of the compositions.

## Conclusion

We have successfully crafted nanoparticles using thiolated arabinoxylan (TAX), incorporating aripiprazole (APZ). The optimized formulation (F4) showcased favourable properties, including a uniform particle size, as revealed by thorough characterizations involving physicochemical aspects, encapsulation efficiency, and in vitro release profiles in simulated gastric fluid (SGF) and simulated intestinal fluid (SIF). Our comprehensive investigation, spanning in vitro techniques, in vivo tests, and histopathological studies, yielded promising results, endorsing the viability and potential of APZ-loaded TAX-based nanoparticles. Toxicity testing confirmed the safety of these nanoparticles, laying the groundwork for their further exploration in in vivo studies. In conclusion, the TAX-based nanoparticle system emerges as a promising avenue for the oral delivery of antipsychotic drugs. This research contributes significantly to drug delivery systems, offering potential advancements in the effective administration of antipsychotic medications.

## Ethics Statements

Guidelines for care and use of laboratory animals of Faculty of Pharmacy, University of Sargodha, Sargodha, Pakistan, were used to perform animal studies, and these studies were duly approved by the animal ethics committee of University of Sargodha.

## Data Availability Statement

The authors confirm that the data supporting the findings of this study are available within the article.

## Author Contributions

Mehwish Sikandar: Methodology, Project administration, Writing—original draft, Software. Ume Ruqia Tulain; Resources, Supervision, Validation, Formal analysis. Nadia Shamshad Malik; Resources, Supervision, Validation, Formal analysis. Alia Erum; Conceptualization, Data curation, Formal analysis. Arshad Mahmood; Conceptualization, Data curation, Resources. Muhammad Tariq Khan; Visualization, Writing—original draft, Conceptualization. Asif Safdar; Visualization, Data curation, Investigation.

